# Context-dependent genetic regulation of gene expression in pigs

**DOI:** 10.64898/2026.03.11.710275

**Authors:** Fan Wang, Chen Wang, Jinyan Teng, Lingzhao Fang, Iuliana Ionita-Laza

## Abstract

Production livestock provide a natural system for studying gene regulation under physiologically demanding conditions shaped by rapid growth, environmental exposure, and immune challenges. Using farm pigs from the PigGTEx resource as a model, we applied quantile regression to uncover latent, context-dependent genetic effects on gene expression across tissues. This approach identifies quantile-specific expression quantitative trait loci (eQTLs) that are not detected by standard linear regression and are enriched in distal regulatory elements and three-dimensional genome architectural features rather than promoter-proximal regions. Genes with quantile-dependent eQTLs are more intolerant to loss-of-function variants and exhibit stronger enrichment in GO functional categories, indicating their likely functional significance. Cross-species comparisons reveal substantial overlap between pig and human eGenes across tissues, indicating conservation of regulatory architecture. Notably, many quantile-specific eQTLs influence tail expression states and involve genes relevant to human disease. For example, we identify a cis-eQTL affecting the conserved transcriptional regulator *BCL6B* in pig blood that modulates enhancer activity and reduces expression at lower quantiles. In contrast, *BCL6B* is minimally expressed in resting human blood and lacks detectable cis-regulatory variation under baseline conditions, consistent with its reported induction during immune activation. These findings demonstrate that pig eQTL maps can reveal context-dependent regulatory variation at loci that remain silent or weakly variable in human cohorts.

## Introduction

Expression quantitative trait loci (eQTLs) link genetic variants to changes in gene expression. Numerous initiatives such as Genotype-Tissue Expression (GTEx) and extended GTEx (eGTEx) in humans as well as Farm animal Genotype-Tissue Expression (FarmGTEx) in livestock support the discovery of causal genetic variants by connecting molecular quantitative trait loci in both humans and animals to relevant diseases^1;2;3^.

FarmGTEx aims to decipher the genetic architecture for gene expression across 16 species under diverse biological and environmental contexts. Since its inception in 2018, FarmGTEx has completed several milestones in cattle, pigs, and chickens using public datasets^4;5;6^. These data include GWAS and transcriptomic data, as well as epigenomic data (histone marks, chromatin accessibility) across multiple tissues etc. Unlike laboratory animals typically used as surrogate organism-level models, farmed animals offer distinct advantages for studying context-specific regulatory effects in natural populations exposed to diverse environmental conditions (for example, diet, climate and pathogen exposure). In this study, we focus on genetic effects on gene expression in pigs (PigGTEx^5^). Pigs serve as a valuable biomedical model for humans due to their physiological, anatomical, and developmental similarities^7^. They share human-like organ sizes, cardiovascular physiology, metabolism, and immune system features, making them relevant for studies across multiple organ systems. In the brain, pig size, organization, myelination, and neurodevelopmental timing are more similar to humans than to rodents, supporting their use in neuroscience research.

Farm pigs live in environments that impose sustained physiological and immunological demands, including high metabolic load, dense housing, pathogen exposure, and production-driven selection for rapid growth and reproduction. As a result, many regulatory pathways in pigs, particularly those related to immunity, stress response, metabolism, and inflammation, are constitutively more active than in humans under baseline conditions. This environmental priming is further evidenced by studies comparing conventionally raised pigs to those in specific pathogen free (SPF) facilities, where SPF pigs exhibit a significantly more quiescent immune profile that aligns more closely with the resting baselines observed in human GTEx data^8^. This “pre-activated” physiological state makes pigs a powerful system for eQTL mapping, as regulatory variants that may be silent or weak in humans can manifest measurable transcriptional effects in pigs without experimental stimulation. eQTL studies in pigs therefore are particularly valuable for uncovering context-dependent regulatory variation, identify genetic control of gene expression under chronic activation, and gain insight into gene-environment interactions that are difficult to capture in human cohorts.

Unlike previous analyses of these data, our approach leverages quantile regression (QR^9^), an extension of linear regression (LR) for quantitative traits that models conditional quantiles of a phenotype rather than only its conditional mean, therefore complementing recent work on these data^5^. This framework allows the detection and characterization of heterogeneous eQTLs, where genetic effects differ across the distribution of gene expression. Whereas LR assumes that genetic effects are constant across the entire distribution of a trait, QR allows genetic effects to vary depending on the level of the phenotype. In particular, causal variants may have only small effects in most healthy individuals (and thus minimal impact on the trait mean) but exert stronger effects in the extremes or tails of a biomarker distribution, where clinically relevant subpopulations are often concentrated. Such associations are often missed by standard LR approaches commonly used in GWAS.

Detecting heterogeneous eQTLs can provide important biological insights, as such variability may reflect gene-by-context interactions or regulatory processes that modulate gene expression across cellular or environmental states^9^. For instance, eQTL effect sizes may vary with factors such as cell type composition, developmental stage, experimental or physiological stimulation, environmental conditions, or disease state, contexts that are not captured by standard LR models, which focus solely on mean effects and assume homogeneous effects across contexts. In contrast, QR is well-suited to uncover latent context-dependent associations, providing a powerful framework for identifying regulatory variants with variable effects and prioritizing genes with potential disease relevance. Increasingly, studies highlight the role of context-specific effects, e.g. response eQTLs, that can help explain disease-associated loci lacking detectable steady-state eQTLs^10^. The key feature of QR is that it can detect such interaction effects without having the context explicitly included in the model. This is indeed an important advantage as the number of possible contexts is very large and cannot be systematically assessed in practice.

Here, we apply quantile regression to five tissues in PigGTEx to characterize eQTL and eGene discoveries, evaluate their functional relevance, and quantify quantile-specific cis-eQTL genetic effects across the gene expression distribution at selected loci implicated in human diseases.

## Results

### PigGTEx expression data

The pilot phase of PigGTEx project^5^ has generated protein-coding gene (PCG) expression data for samples from several tissues. We focused on expression data in five tissues with the largest sample sizes (muscle *n* = 1, 321; liver *n* = 501; brain *n* = 419; blood *n* = 386; adipose *n* = 285) and used the preprocessed expression matrices, genotype data, and covariates made available by Teng et al.^5^. Briefly, for cis-eQTL mapping, we considered only PCG with TPM ≥ 0.1 and/or raw read counts ≥ 6 in at least 20% of samples. PCG expression was normalized across samples within each tissue using the trimmed mean of M-values (TMM) method implemented in edgeR^11^, followed by inverse normal transformation. The genotype dataset included 3, 087, 268 SNPs, filtered for a minor allele frequency (MAF) greater than 0.05 within each tissue. To account for genetic structure and technical confounding, we adjusted for the top ten principal components of genetic variation and the top ten PEER factors estimated from PEER (v1.0) R package in our analyses^5^.

### Comparing baseline immune activation in pigs vs. humans

We examined representative genes from key immune pathways, including *OAS1, ISG15, IFNG, MX1, IL2RA, TNF*, and *CXCL10* to assess baseline immune activation in pigs vs. humans. We observed notable differences between pig samples in PigGTEx and human GTEx (v10) data (**Fig. 1**), with pigs showing higher expression of several genes in whole blood, consistent with a partially active or chronically stimulated antiviral state. In contrast, human samples exhibited the expected low basal expression of these genes in healthy, unchallenged individuals. This pattern is consistent with prior studies reporting elevated baseline immune activity in farm-raised pigs relative to animals maintained under controlled laboratory or high-biosecurity conditions^8^, suggesting that environmental exposure and production conditions contribute to a more immune-primed physiological state in production livestock.

**Figure 1:**
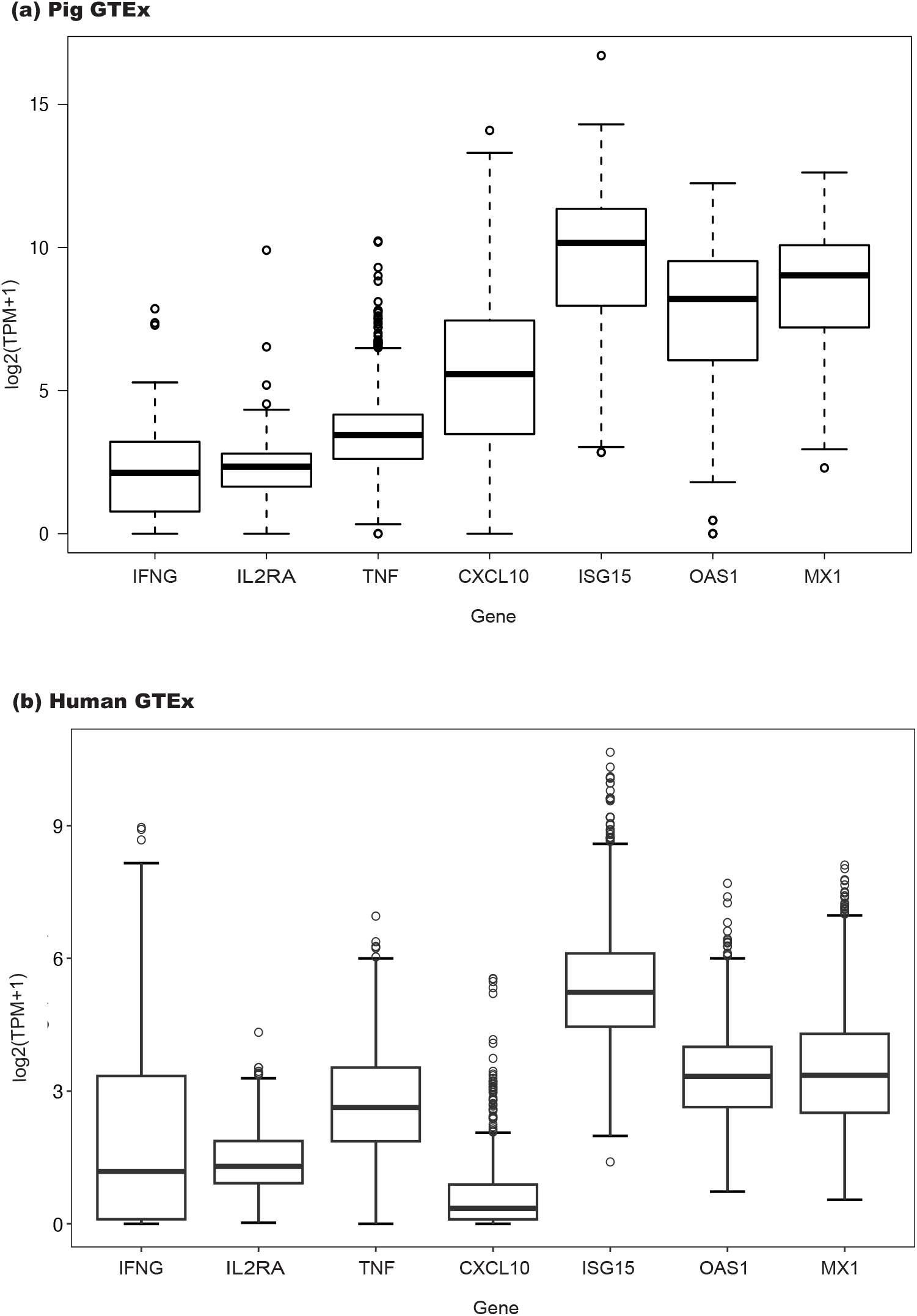
Comparative expression of immune-related genes in farm pigs in PigGTEx vs. human GTEx baseline in blood. Boxplots represent the distribution of transcript abundance (log2(TPM+1)) for key cytokines and interferon-stimulated genes.

### eQTL and eGene discovery across five tissues

To account for polygenic effects and population genetic structure, we used recent genome-wide regression methods including Regenie^12^ for LR and Regenie.QRS^13^ for QR (**Methods**). Using both methods, we performed cis-eQTL mapping within a cis window defined as *±*1Mb around the transcription start site (TSS) of each PCG. For both Regenie and Regenie.QRS, and for a given PCG, we first generated polygenic predictions for all individuals using ~ 10, 000 pruned genome-wide SNPs based on *r*^2^ = 0.2. For Regenie.QRS, the quantile rank score test was applied at a grid of quantile levels *τ* = (0.1, …, 0.9) and quantile-level *p*-values were combined using the Cauchy combination method^14^. To assess gene-level significance, we used again the Cauchy combination method to aggregate *p*-values across SNPs for each gene, followed by Benjamini-Hochberg (BH) false discovery rate (FDR) control at the 0.05 level across all genes. The significant genes are referred to as eGenes.

Using Regenie.QRS, we detected 8,023 (59.63%) eGenes in muscle, 4,958 (37.44%) in liver, 4,519 (35.78%) in blood, 3,781 (24.94%) in brain and 3,122 (21.73%) in adipose. Corresponding eGene counts obtained with Regenie were similar and are reported in **Table 1**. We additionally report, for each tissue and method, the number of significant cis-eQTLs (SNPs with *p* < 1 × 10^−6^).

**Table 1:** Number of eGenes and eQTL discoveries. Number of eGenes (FDR < 0.05) and their proportion among PCG per tissue for each method are reported. Also shown are the number of significant cis-eQTLs (*p* < 1 × 10^−6^) for each tissue and method.

| Tissue | PCG | Regenie |  | Regenie.QRS |  |
| --- | --- | --- | --- | --- | --- |
|  |  | eGenes | eQTLs | eGenes | eQTLs |
| Muscle | 13,454 | 8,426 (62.63%) | 1,406,803 | 8,023 (59.63%) | 1,213,038 |
| Liver | 13,243 | 5,119 (38.65%) | 850,257 | 4,958 (37.44%) | 698,448 |
| Blood | 12,629 | 4,526 (35.84%) | 673,771 | 4,519 (35.78%) | 549,210 |
| Brain | 15,161 | 4,033 (26.60%) | 551,315 | 3,781 (24.94%) | 422,064 |
| Adipose | 14,364 | 3,059 (21.30%) | 276,959 | 3,122 (21.73%) | 183,905 |

Although most eGenes identified by Regenie and Regenie.QRS are shared (**Fig. 2**), Regenie.QRS yields a substantial number of additional discoveries that are missed by Regenie, suggesting potential heterogeneity in gene expression regulation. Notably, these additional eGenes are significantly enriched in Gene Ontology (GO) functional categories, to a greater extent than eGenes identified only by Regenie (**Methods, Fig. S1-S10**). Notably, enriched categories include biological processes related to cellular responses to stress, chemical stimuli, and endogenous signals, consistent with the idea that these genes can be regulated by context-dependent eQTLs that are only active under specific stimulations. We also observe considerable sharing of eGenes across tissues, including many genes shared across all five tissues; however, many eGenes are also only detected in a single tissue (**Fig. S11**). This pattern partly reflects genuine tissue-specific regulation, but may also arise from weak shared signals that are difficult to detect consistently across tissues and methods due to limited statistical power.

**Figure 2:**
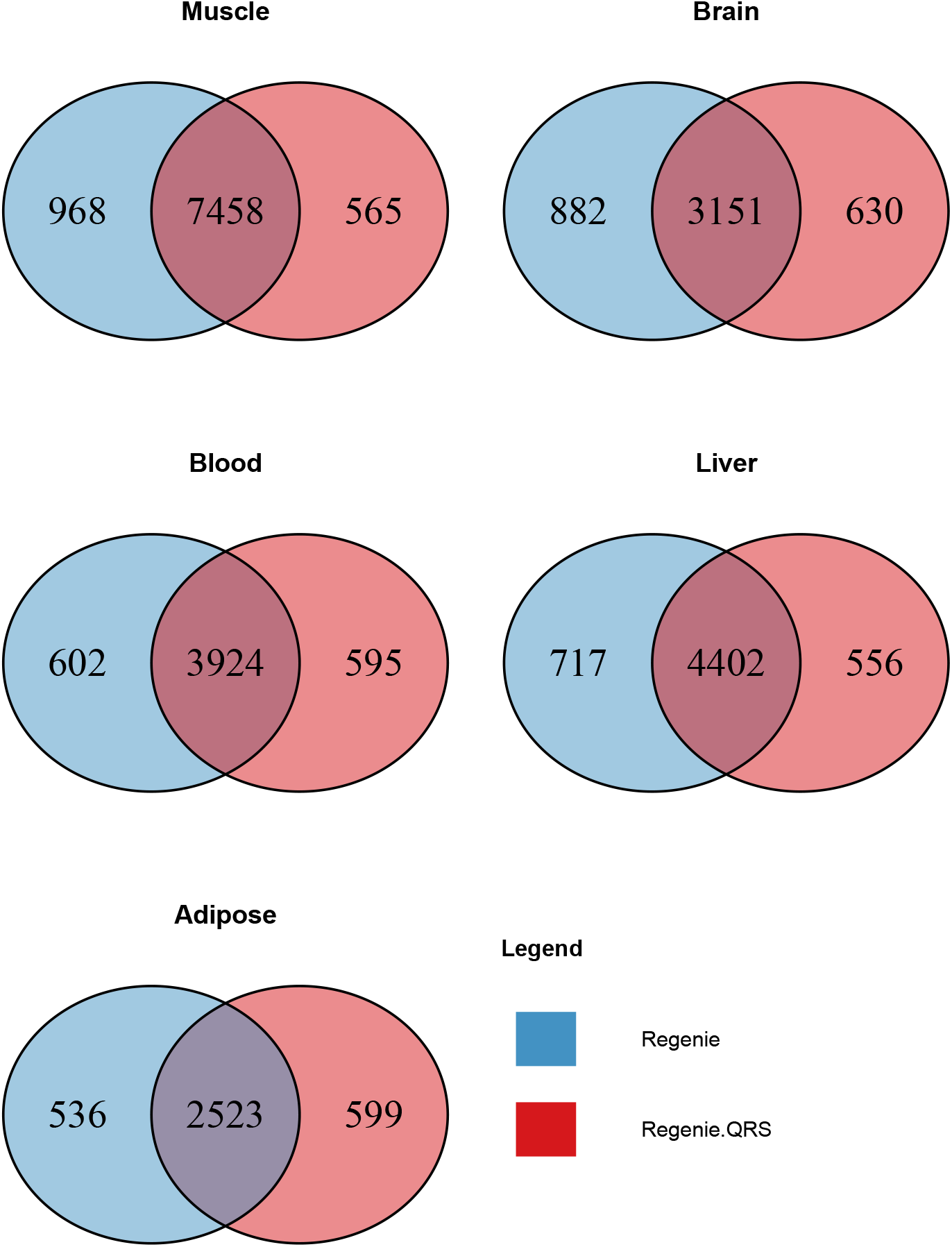
Overlap of eGenes identified by Regenie and Regenie.QRS across five tissues.

We next assessed the overlap between pig and human eGenes identified by GTEx (v10) in matched tissues. Pig eGenes were first mapped to their one-to-one human orthologs, and genes without a clear orthologue were excluded. The resulting set of human orthologues (~ 90% of the original pig eGenes) was then compared with GTEx eGenes to quantify cross-species overlap. Overall, a substantial fraction of pig eGenes is also detected in GTEx, with similar overlap proportions across methods (**Fig. S12**). In particular, brain exhibits the highest proportion of shared eGenes, consistent with the well-documented regulatory and functional similarity between pig and human brain. Notably, liver shows a substantially lower degree of overlap compared with other tissues. This reflects in part the relatively limited sample size of human liver GTEx data, which results in fewer detected human liver eGenes.

### Regenie.QRS eQTLs show heterogeneous effects across the expression distribution

Because QR enables estimation of effects at multiple quantiles, it allows us to assess the degree of heterogeneity in eQTL effects across the distribution. For each eQTL, we define a heterogeneity index as the log-transformed ratio of the standard deviation to the absolute mean of the estimated quantile coefficients: log(sd(*β*(*τ*))*/*|mean(*β*(*τ*))|. A higher index indicates greater variability in effect sizes across quantiles. Consistent with this, eQTLs detected by Regenie.QRS but missed by Regenie show the highest heterogeneity (**Fig. 3(a)**).

**Figure 3:**
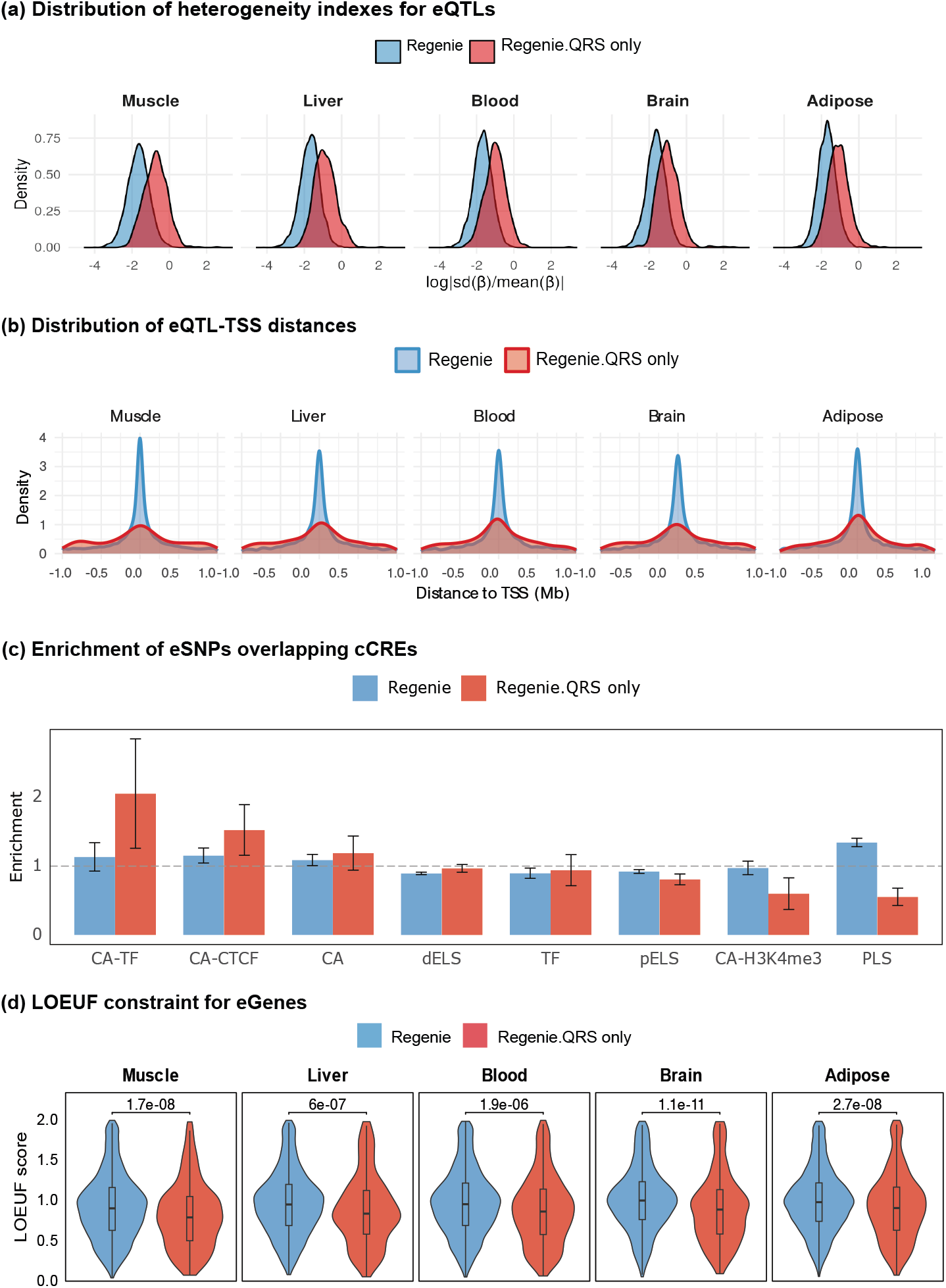
eQTL heterogeneity, distance to TSS, regulatory activity and eGene constraint. (a) Densities of heterogeneity indexes for eQTLs detected by Regenie and Regenie.QRS only, (b) Distribution of distances of lead eQTL from Regenie and Regenie.QRS only to the TSS of eGenes, (c) Enrichment of eSNPs within cCREs and standard errors, (d) Comparison of gene constraint (LOEUF) distributions for eGenes detected by Regenie vs. Regenie.QRS only across tissues.

Moreover, the proportion of positive eQTL effects decreases monotonically across quantiles, with a higher proportion (≥ 0.5) of positive effects at low quantiles and a lower proportion (≤ 0.5) at high quantiles (**Fig. S13**). The decrease is steepest for Regenie.QRS only eQTLs, in line with the increased heterogeneity index above. This pattern is consistent across the five tissues and suggests that a greater proportion of alleles tend to increase gene expression in the lower tail of the distribution, whereas a greater proportion tend to decrease expression in the upper tail. One possible interpretation of this quantile-dependent pattern is the presence of biological floor or ceiling constraints on gene expression, whereby allelic effects are more likely to increase expression in low-expression states but attenuate or reverse in high-expression states, consistent with regulatory saturation or homeostatic buffering mechanisms.

### Regenie.QRS captures eQTLs beyond promoter-proximal regions

For each eGene, we computed the distance between the lead eQTL and the gene TSS. For each tissue, we compared the resulting distance distributions between two discovery categories: eGenes identified by Regenie and eGenes identified exclusively by Regenie.QRS. Across tissues, eQTLs associated with eGenes detected by Regenie were more strongly concentrated near the TSS, whereas those detected by Regenie.QRS exhibited broader distance distributions with heavier tails. This pattern suggests that quantile-specific associations are, on average, less tightly localized to promoter-proximal regions and may more frequently involve distal regulatory elements such as enhancers (**Fig. 3(b)**).

To further validate our findings, we performed enrichment analyses of eSNPs within candidate Cis-Regulatory Elements (cCREs) annotated by ENCODE4^15^. For each of the 28,108 eGenes identified by Regenie and Regenie.QRS across five tissues in PigGTEx, we defined the corresponding lead eQTL as the representative eSNP. Of these, 15,535 pig eSNPs were successfully lifted over to the human reference genome (hg38) using UCSC liftOver^16^.

Enrichment was quantified by comparing the proportion of eSNPs overlapping cCREs to that of matched random SNPs. For each eSNP, we generated 50 control gene-SNP pairs matched on the distance between the SNP and the gene TSS, using greedy nearest-neighbor matching implemented in the MatchIt R package^17^. Matching on TSS distance accounts for the strong positional bias in regulatory element distribution relative to genes. This step is particularly important because eQTLs identified by Regenie.QRS tend to lie farther from gene TSS than those detected by Regenie, and failing to control for this difference could confound enrichment estimates. Enrichment was evaluated using odds ratios (ORs) derived from Fisher’s exact test based on 2 × 2 contingency tables. We evaluated several functional categories, including chromatin-accessible regions with strong transcription factor binding (CA-TF), chromatin-accessible regions with strong CTCF binding sites (CA-CTCF), distal enhancer-like elements (dELS), and promoter-like sequences (PLS). Compared to standard Regenie eQTLs, SNPs uniquely identified by Regenie.QRS exhibited significantly higher enrichment in CA-TF and CA-CTCF categories, while displaying a pronounced depletion in promoter-proximal elements (**Fig. 3(c)**).

### Regenie.QRS detects eQTLs with stronger AlphaGenome-predicted regulatory effects

We also used AlphaGenome^18^ to predict the regulatory impact of eQTLs identified by Regenie and Regenie.QRS. AlphaGenome estimates variant effects using 5,930 human omics tracks, which can be grouped into 11 molecular modalities (**Table S1**), and outputs one score per track for each variant. Within each modality, we summarized predictions by taking the maximum score across all corresponding tracks. Approximately 1.93 million pig eQTLs with Regenie and/or Regenie.QRS *p* < 10^−6^ were lifted over to human genomic coordinates, yielding approximately 975, 483 variants. Note that only genomic coordinates were lifted over, while the REF/ALT alleles from the pig variants were retained. In addition to the 11 modality-specific scores, we also derived a global score by taking the maximum score across all modalities.

#### Tissue- and gene-agnostic multi-modal functional prediction

We compared functional predictions for eQTLs identified by Regenie and Regenie.QRS, considering all AlphaGenome scores irrespective of tissue context or the target gene of the human omics track. Across five PigGTEx tissues and twelve AlphaGenome modalities, a total of 60 comparisons were performed, leading us to adopt a Bonferroni-corrected significance threshold of 0.05*/*60 = 0.0008. Regenie.QRS eQTLs exhibited significantly higher functional scores than Regenie eQTLs for several modalities such as RNA-seq, Splice sites and Chromatin contact maps in brain (**Table 2**). By contrast, Regenie eQTLs only showed significantly higher scores for the global AlphaGenome score in liver.

**Table 2:**
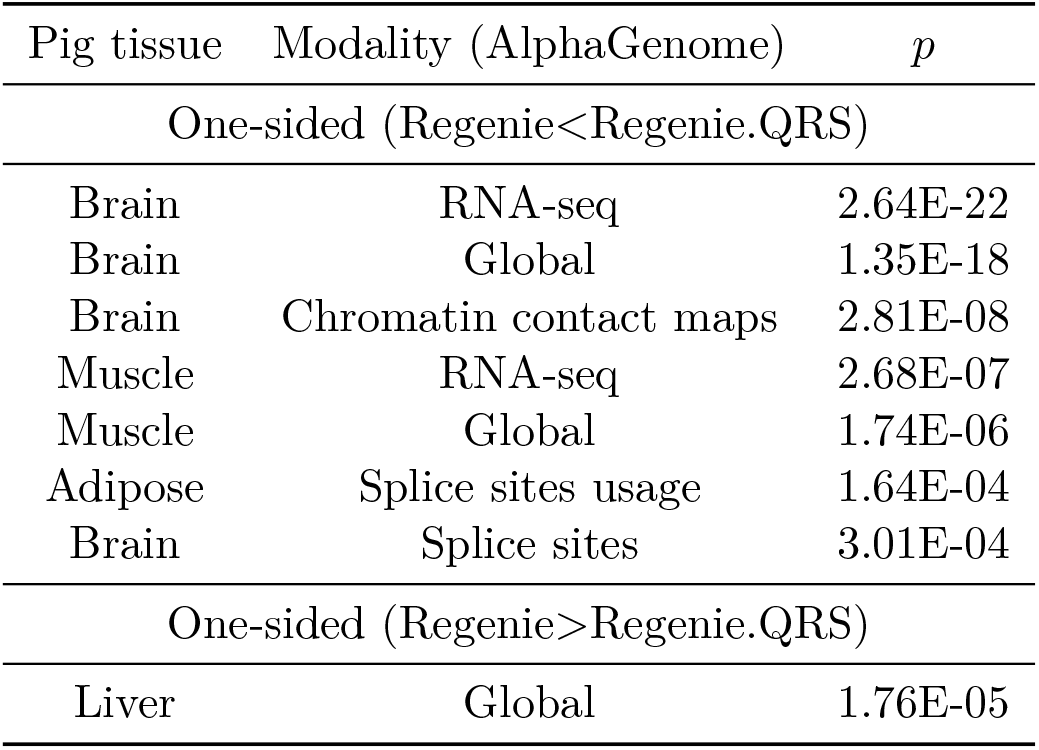
AlphaGenome prediction of eQTL effects. Comparison of AlphaGenome scores for eQTLs identified by Regenie and Regenie.QRS. *p*-values are from one-sided Wilcoxon test. Results are shown for tissue- and gene-agnostic multi-modal annotations.

#### Tissue-matched regulatory effects independent of target gene

We then restricted comparisons to eQTLs for which the tissue context of the human gene expression regulation track matched the PigGTEx tissue, without requiring that the target gene correspond to the pig eGene. Only gene expression regulation modality scores were considered, as their tissue definitions align well with PigGTEx tissues. Within this tissue-matched framework, Regenie.QRS eQTLs exhibited significantly higher scores than Regenie eQTLs for brain and muscle, whereas no tissue showed significantly higher scores for Regenie compared with Regenie.QRS (**Table S2**).

#### Tissue- and gene-matched regulatory effects

Finally, we restricted comparisons to eQTLs for which both the tissue context and the target gene of the human gene expression regulation track matched the PigGTEx tissue and eGene. In this fully matched setting, Regenie.QRS eQTLs showed significantly higher functional scores than Regenie eQTLs in brain, whereas Regenie eQTLs exhibited significantly higher scores in adipose and blood (**Table S3**).

### Regenie.QRS eGenes show higher loss-of-function constraint

We evaluated whether eGenes identified only by Regenie.QRS are more intolerant to loss-of-function (LoF) variation compared to eGenes identified by Regenie. We used LOEUF (Loss-of-function Observed/Expected Upper bound Fraction), a gene-level constraint metric developed by the gnomAD project that quantifies a gene’s tolerance to LoF mutations in the human population. Across all five tissues analyzed, Regenie.QRS eGenes exhibited significantly lower LOEUF scores than Regenie eGenes, indicating greater intolerance to LoF variation (**Fig. 3(d)**). This result supports the hypothesis that heterogeneous eQTL effects that are potentially context-dependent are more likely to occur in genes under stronger selective constraint^10^, reinforcing the potential relevance to disease of eGenes identified by Regenie.QRS.

### Pig eQTLs help identify disease relevant human genes

We next evaluated the extent of overlap between pig eGenes and genes listed in the GWAS Catalog. A pig eGene was defined as overlapping if its corresponding one-to-one human ortholog was found among GWAS Catalog genes with genome-wide significant associations. The overlap proportion was calculated as the number of overlapping genes divided by the total number of pig eGenes. Across all five tissues, eGenes uniquely identified by Regenie.QRS consistently showed a high overlap with GWAS Catalog genes (over 60%), and this proportion was slightly higher than that for eGenes identified by Regenie (**Fig. S14**).

Together, these results support the functional relevance and human translatability of eQTLs and eGenes identified by Regenie.QRS. We next highlight several loci with heterogeneous effects across quantiles detected by Regenie.QRS but missed by Regenie and that map to known human disease loci. For each locus, we show cis-eQTL association results from Regenie and Regenie.QRS as well as quantile-specific fine-mapping results using QR-CARMA (**Methods**), focusing on the specific quantile that produced a credible set at each locus.

#### *BCL6B* and blood

*BCL6B* (B-cell lymphoma 6B) is a highly conserved transcriptional regulator involved in immune cell activation and acts as a tumor suppressor in multiple cancers^19;20^. In humans, *BCL6B* expression is low or absent in resting lymphocytes but is induced upon immune activation and consequently detectable in a subset of antigen-experienced CD8^+^ T cells^21^, which likely explains the absence of detectable cis-eQTLs for this gene in whole-blood GTEx analyses, even when using quantile-based approaches.

We identified *BCL6B* as an eGene using Regenie.QRS only, with lead cis-regulatory variant (chr12:51,968,848; MAF = 0.09; *p* = 3.3 · 10^−8 &^ PIP = 0.20 at quantile *τ* = 0.1) with heterogeneous effects across expression quantiles (**Fig. 4**). Notably, the genetic effect is strongest and negative at lower expression quantiles, while the effect diminishes at higher quantiles. This eQTL also affects the expression of a blood enhancer (chr12:51,975,801-51,976,200; 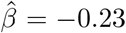, *p* = 10^−5^), situated within a highly conserved region and overlaping established human regulatory elements, including ENCODE4 cCREs and GeneHancer (**Fig. 4**). *BCL6B* exhibits a significant, progressive increase in expression in porcine blood from fetal to adult stages (**Fig. S15**, Pig developmental GTEx project), consistent with developmental changes in the immune system. Expression levels do not differ between males and females (**Fig. S15**).

**Figure 4:**
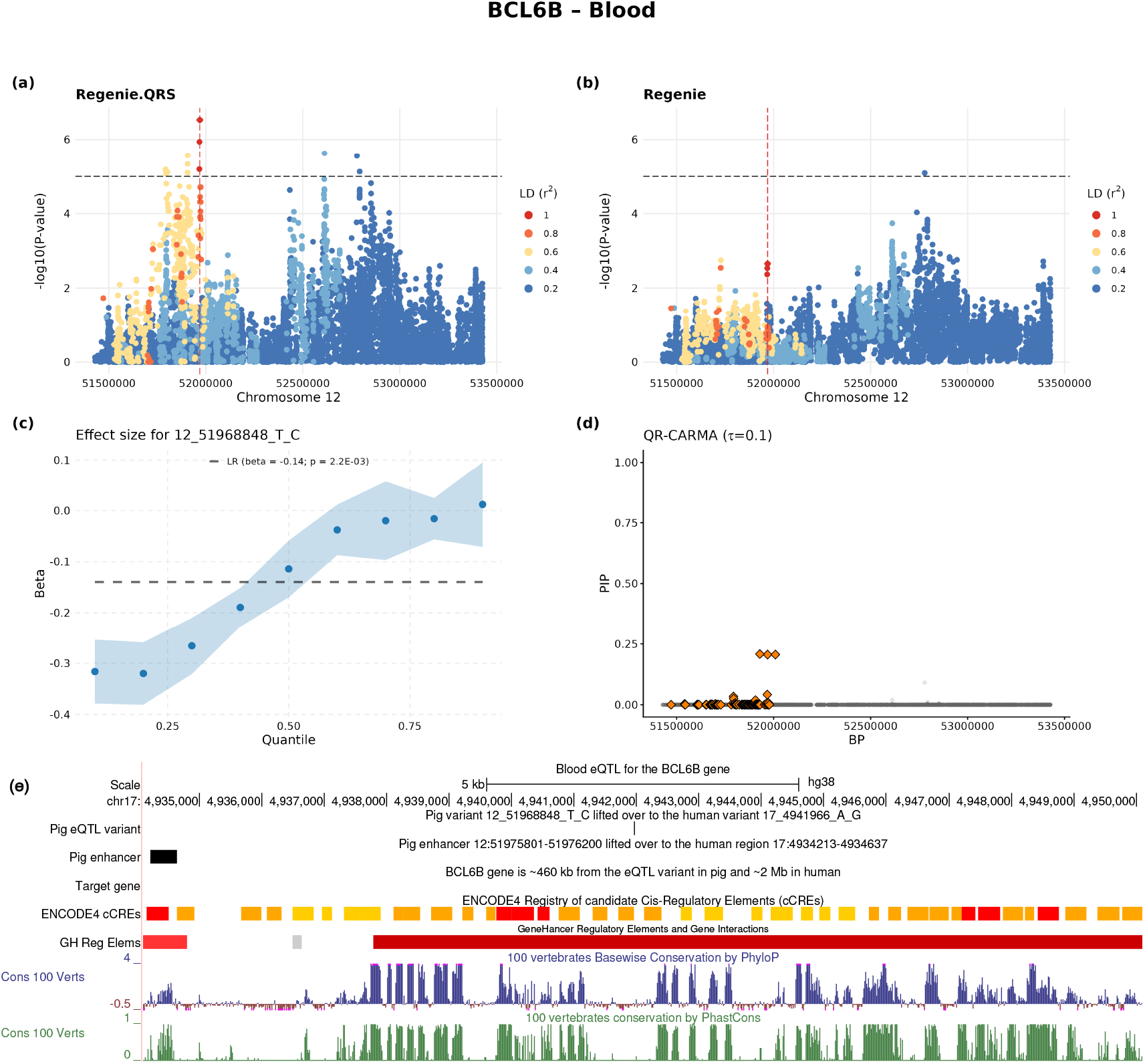
Heterogeneous eQTL effect at *BCL6B* gene (ENSSSCG00000017927) in blood tissue. (a–b) LocusZoom plots show − log_10_(*p*-value) for cis-eQTL associations from (a) Regenie.QRS and (b) Regenie, with the lead Regenie.QRS eQTL indicated by a diamond and a vertical dashed line. Point colors represent linkage disequilibrium (LD; *r*^2^) with the lead variant. The horizontal dashed line marks the significance threshold at 10^−5^. (c) Quantile-specific effect sizes (*±* standard errors) for the lead variant across quantile levels, with the LR estimate shown as a dashed line. (d) QR-CARMA fine-mapping results at quantile *τ* = 0.1, showing PIPs for variants across the locus; SNPs in credible sets (*ρ* = 0.9, *r* = 0.6) are highlighted. (e) Genomic context of the top PIP eQTLs and the associated enhancer. The data is mapped to the human hg38 genome assembly on chromosome 17. The enhancer is situated within a highly conserved region and overlaps established human regulatory elements (ENCODE4 cCREs and GeneHancer).

Although *BCL6B* lacks detectable steady-state eQTLs in human blood, the presence of a functional, quantile-dependent cis-eQTL in pig suggests that *BCL6B* expression can be genetically regulated in blood immune cells and is consistent with a model in which genetic variation at this locus may influence immune-related traits in humans in an activation-specific manner.

#### *ITGA5* and muscle

*ITGA5* (Integrin Subunit Alpha 5) encodes the *α*5 subunit of the fibronectin receptor *α*5*β*1 integrin and contributes to muscle development and tissue homeostasis by mediating cell-extracellular matrix adhesion and mechanotransduction^22^. In mesenchymal stromal cells (MSCs), *ITGA5* plays a well-established role in promoting osteoblast differentiation^23^. In skeletal muscle, *ITGA5*-fibronectin signaling is critical for muscle fiber organization, regeneration, and response to mechanical load^24^.

We identified *ITGA5* as an eGene in muscle using Regenie.QRS only, with lead eQTL (chr5:18,693,282; MAF = 0.05; *p* = 2.1 · 10^−8 &^ PIP = 0.40 at quantile *τ* = 0.1) with significant negative effect among individuals with lower expression, and increasing effects across quantiles (**Fig. 5**). This eQTL also affects the expression of a pig muscle enhancer (chr5:18,436,001-18,437,600; 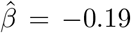, *p* = 1.29 · 10^−4^) situated in a highly conserved region, and overlaping established human regulatory elements, including ENCODE4 cCREs and GeneHancer (**Fig. 5**).

**Figure 5:**
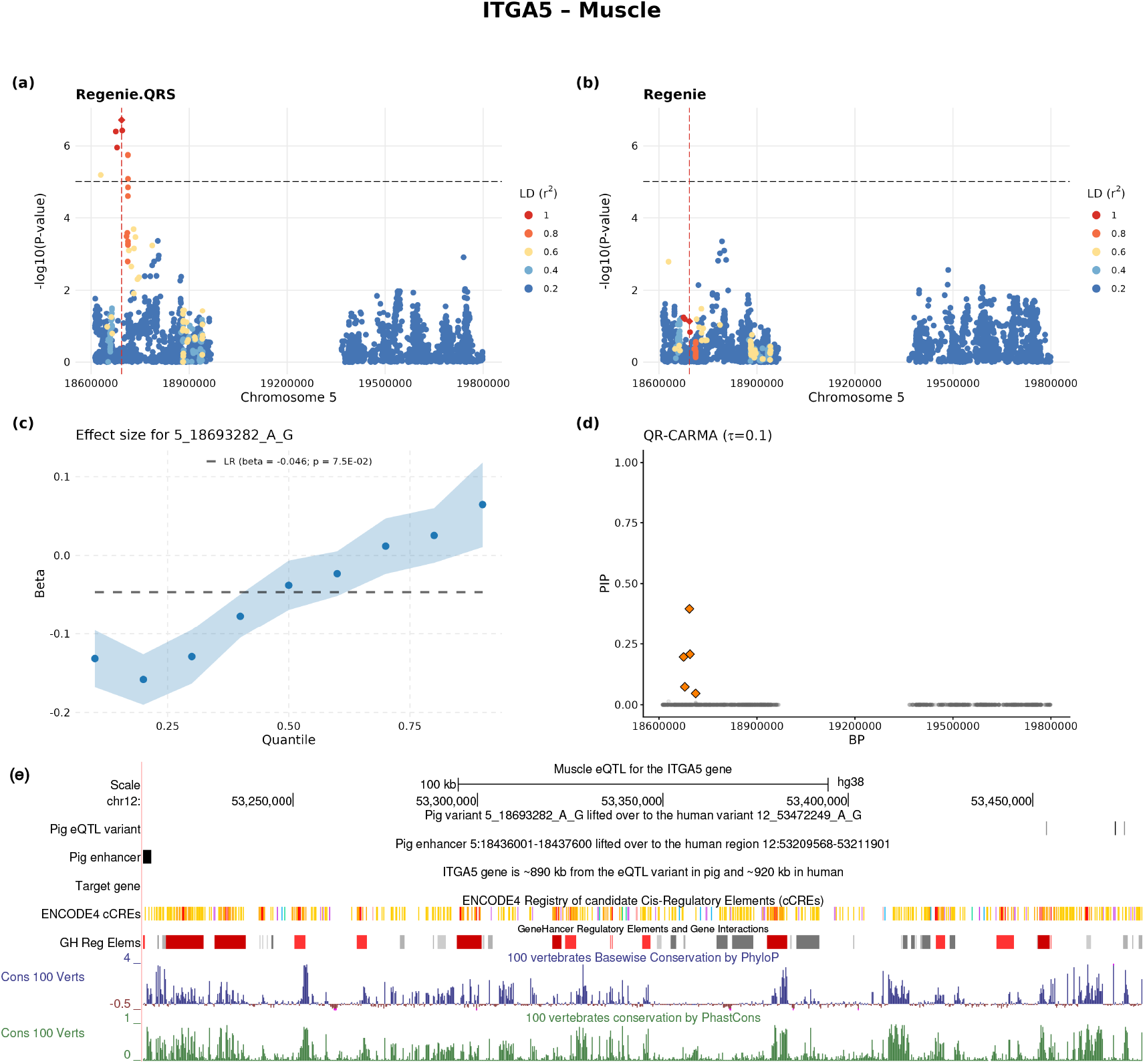
Heterogeneous eQTL effect at *ITGA5* gene (ENSSSCG00000000293) in muscle tissue. (a–b) LocusZoom plots show − log_10_(*p*-value) for cis-eQTL associations from (a) Regenie.QRS and (b) Regenie, with the lead Regenie.QRS eQTL indicated by a diamond and a vertical dashed line. Point colors represent linkage disequilibrium (LD; *r*^2^) with the lead variant. The horizontal dashed line marks the significance threshold at 10^−5^. (c) Quantile-specific effect sizes (*±* standard errors) for the lead variant across quantile levels, with the LR estimate shown as a dashed line. (d) QR-CARMA fine-mapping results at quantile *τ* = 0.1, showing PIPs for variants across the locus; SNPs in credible sets (*ρ* = 0.95, *r* = 0.5) are highlighted. (e) Genomic context of the top PIP eQTLs and the associated enhancer. The data is mapped to the human hg38 genome assembly on chromosome 12. The enhancer is situated within a highly conserved region and overlaps established human regulatory elements (ENCODE4 cCREs and GeneHancer).

These findings suggest that variation at the *ITGA5* locus could contribute to muscle-related traits and disease susceptibility through effects on cell-matrix interactions and tissue remodeling.

#### *GFAP* and brain

*GFAP* (Glial Fibrillary Acidic Protein) is a core astrocyte cytoskeletal gene whose expression increases during reactive astrogliosis following diverse CNS perturbations^25;26^. Heterozygous gain-of-function mutations cause Alexander disease, a monogenic leukodystrophy with severe neurodevelopmental and neurodegenerative features. Additionally, common and low-frequency *GFAP* variants associate with white-matter pathology, in broader cerebrovascular and aging-related brain disease^27;28;29^.

We detected *GFAP* as an eGene using Regenie.QRS but not Regenie. The lead eQTL variant at the locus (chr12:18,504,738; MAF = 0.1; *p* = 1.4 · 10^−7 &^ PIP = 0.18 at quantile *τ* = 0.1) shows clear effect size heterogeneity across the expression distribution, with negative effects at lower quantiles and positive effects toward the upper quantiles (**Fig. 6**). This gene is highly expressed in both pig and human brain tissue, with significant upregulation during the postnatal period compared to the prenatal stage^30^ (**Fig. S16**).

**Figure 6:**
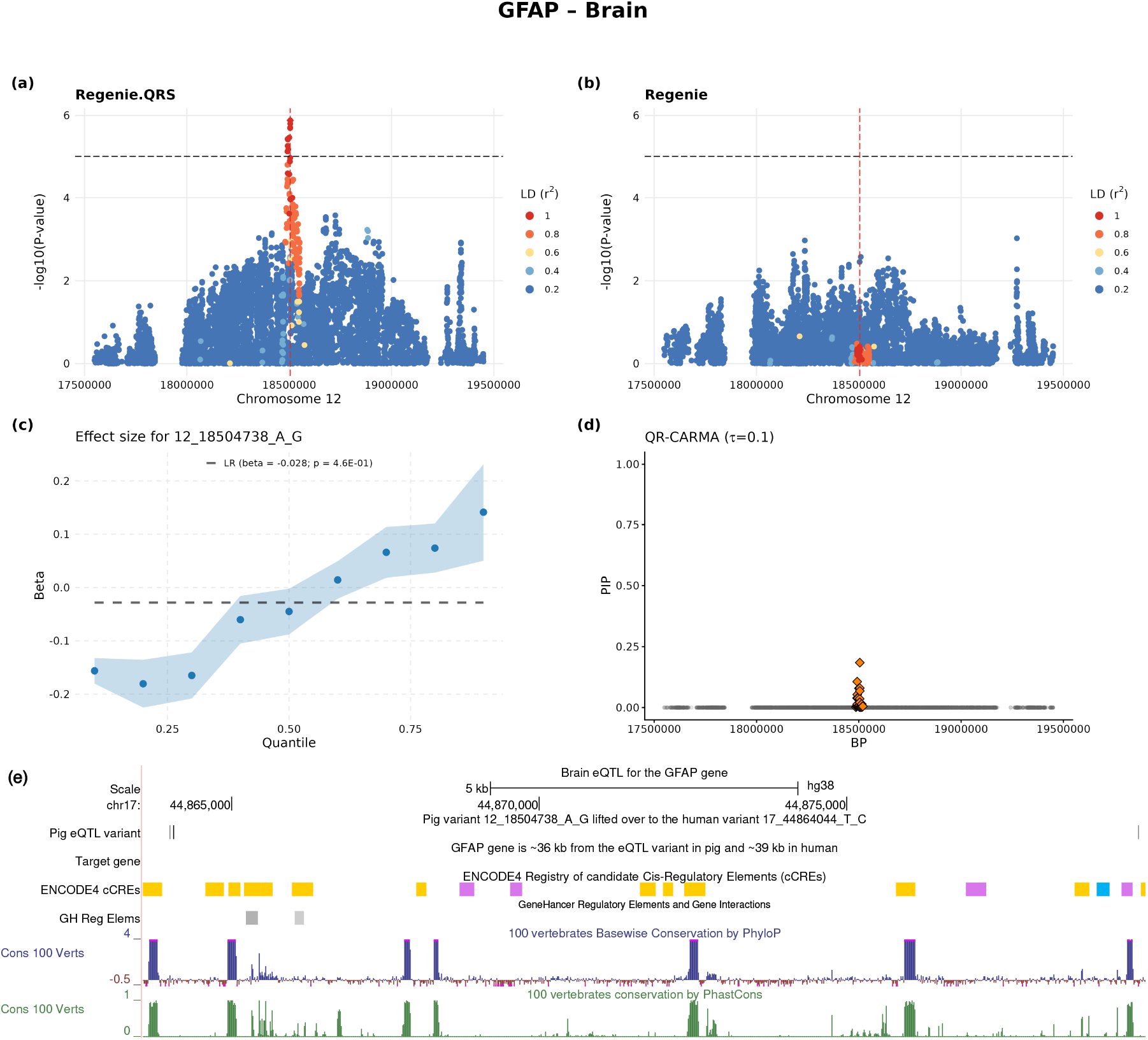
Heterogeneous eQTL effect at *GFAP* gene (ENSSSCG00000017343) in brain tissue. (a–b) LocusZoom plots show − log_10_(*p*-value) for cis-eQTL associations from (a) Regenie.QRS and (b) Regenie, with the lead Regenie.QRS eQTL indicated by a diamond and a vertical dashed line. Point colors represent linkage disequilibrium (LD; *r*^2^) with the lead variant. The horizontal dashed line marks the significance threshold at 10^−5^. (c) Quantile-specific effect sizes (*±* standard errors) for the lead variant across quantile levels, with the LR estimate shown as a dashed line. (d) QR-CARMA fine-mapping results at quantile *τ* = 0.1, showing PIPs for variants across the locus; SNPs in credible sets (*ρ* = 0.95, *r* = 0.5) are highlighted. (e) Genomic context of the top PIP eQTLs. The data is mapped to the human hg38 genome assembly on chromosome 17.

## Discussion

We present an application of QR for the detection of context-dependent eQTLs in pigs, a system with intrinsic advantages arising from natural exposure to heterogeneous environmental conditions, including variation in diet, climate, and pathogen exposure. We show that eQTLs identified in QR tend to have more heterogeneous effects at different quantiles of the gene expression distribution relative to eQTLs identified in LR. Furthermore, we show that eQTLs captured by QR tend to be less concentrated on the promoter-proximal regions, have stronger regulatory function as predicted by AlphaGenome and overlap with cCREs in ENCODE4. Notably, eGenes corresponding to heterogeneous eQTLs tend to be under more evolutionary constraint than eGenes detected by LR, and show significant enrichment in GO functional categories including categories related to cellular responses to stress, chemical stimuli, and endogenous signals, pointing toward roles in stimulus-responsive regulatory processes. Overall, these results point to QR leading to new discoveries with functional consequences and potential relevance to human disease.

Although context dependency and heterogeneity, including gene-by-gene and gene-by-environment interactions, are thought to be widespread, it is practically impossible to comprehensively quantify all possible contexts. Consequently, many existing studies are limited by data collected in restricted settings, including one environmental context at a time and variants close to genes^10;31;32;33;34^. A key advantage of quantile regression is its ability to detect context-dependent effects without explicitly modeling the underlying interaction or non-additive effects. This way, QR can be used to prioritize variants whose effects may be context-dependent including variants in distal enhancers.

In the current study we focused on pigs and have shown the potential of QR to identify eQTLs in pigs that may only be visible in humans under specific contexts or stimulated conditions. Unlike human cohorts, which typically capture gene expression in relatively quiescent baseline states, pigs raised in production environments experience chronic activation of pathways related to growth, immunity, and stress adaptation. This context can amplify the effects of regulatory variants, enabling the detection of eQTLs that may remain weak or context-dependent in humans. Our results therefore highlight the value of pig eQTL maps as a complementary resource for understanding gene regulation in activated biological states, and suggest that livestock populations can reveal dimensions of regulatory architecture that are difficult to access in human studies without experimental perturbation.

Although quantile regression can indirectly capture context dependency by identifying eQTLs with heterogeneous effects across the gene expression distribution, the underlying context inducing this heterogeneity is not explicitly identified, and targeted experiments under specific conditions will be required to elucidate it. Future developments of PigGTEx will incorporate additional contexts, including sex, developmental stages, and diverse environmental conditions which will further advance our understanding of context-dependent associations. Beyond gene expression, QR is broadly applicable to other molecular traits such as chromatin accessibility, DNA methylation, protein abundance, and metabolite levels.

Our PigGTEx study demonstrates that eQTLs detectable in pigs, including context-dependent effects not observed in baseline human samples, can reveal latent regulatory patterns and prioritize genes and pathways for functional follow-up. This approach leverages evolutionary conservation and animal models to complement human data and strengthen biological inference.

## Acknowledgments

We thank Joseph Gogos and Krzysztof Kiryluk for helpful discussions. This research has been partially supported by NIH grant AG072272 and grant 2024-04735 from the Swedish Research Council (to I.I.L.).

## Methods

### eQTL discovery methods

#### Regenie

Regenie^12^ is a whole-genome regression framework that uses a two-step ridge regression approach to efficiently model polygenic effects and control for population structure in large-scale GWAS. In Step 1, it builds a whole-genome predictor by stacked block ridge regression: the genotype matrix is partitioned into consecutive, non-overlapping SNP blocks, and within each block a small set of ridge-regression predictors is generated across multiple shrinkage settings (“Level 0”). These block predictors are rescaled and then combined using a second ridge regression (“Level 1”), with the ridge parameter selected using *K*-fold cross-validation to control overfitting. In Step 2, each tested variant is evaluated in a simple linear model using phenotype residuals after removing the polygenic effect estimated in Step 1.

#### Regenie.QRS

Regenie.QRS^13^ is an extension of Regenie to the QR framework that combines polygenic effect adjustment with scalable quantile regression. In Step 1, Regenie.QRS estimates *ĝ* using Regenie-style polygenic prediction. In Step 2, it fits standard linear quantile regression to *R* = *Y* − *ĝ* by treating *ĝ* as a fixed offset. For inference at a given quantile *τ*, Regenie.QRS uses the quantile rank score (QRS) test, and under the null, the statistic follows 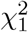 distribution. The asymptotic null does not depend on the phenotype distribution, so it does not require normalizing transformations for validity. In practice, QRS is run across equally spaced quantile levels *τ* = (0.1, 0.2, …, 0.9) and quantile-specific *p*-values are combined using a Cauchy combination to produce a single SNP-level *p*-value.

### g:Profiler for functional enrichment analysis

Functional enrichment analysis was performed using g:Profiler (Sus scrofa version), a web-based tool that integrates multiple biological databases to identify overrepresented functional categories in a gene set. g:Profiler allows for enrichment testing across Gene Ontology (GO) terms for biological processes, molecular functions, and cellular components, as well as pathways from Reactome, KEGG, and other curated resources. We report those GO terms with adjusted p-value less than 0.05.

### QR-CARMA: Quantile-specific fine-mapping with summary-level rank scores

In addition to association testing using Regenie.QRS, at loci of interest we perform statistical fine-mapping using summary statistics from Regenie.QRS obtained at specific quantile level *τ*. We focus on such a locus, and assume there are *p* genetic variants in the region.

Let **R** = (*R*_1_, …, *R*_*N*_)^⊤^ denote the residualized phenotype vector obtained from Regenie.QRS (Step 1), **G** the *N* × *p* genotype matrix with *j*th column **G**_**j**_, and **C** the *N* × *m* matrix of covariates. For a given quantile level *τ*, consider the joint QR model:

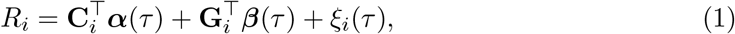

where ***β***(*τ*) = (*β*_1_(*τ*), …, *β*_*p*_(*τ*))^⊤^ is the vector of quantile effects for all SNPs at the locus of interest, and *ξ*_*i*_(*τ*) is the regression error, whose *τ* th conditional quantile given (**G**_*i*_, **C**_*i*_) is zero. We first obtain the QR score statistics and their corresponding *Z*-scores as **S**_*τ*_ = (*S*_1,*τ*_, …, *S*_*p,τ*_)^⊤^ and **Z**_*τ*_ = (*Z*_1,*τ*_, …, *Z*_*p,τ*_)^⊤^. It can be shown that:

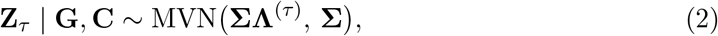

where **Σ** is the LD matrix among the *p* SNPs in the region and **Λ**^(*τ*)^ is the vector of scaled quantile effect sizes. Under the null hypothesis *H*_0_ : ***β***(*τ*) = **0, Z**_*τ*_ ~ MVN(**0, Σ**); under local alternatives, the mean is linear in the underlying quantile effects ***β***(*τ*), allowing the QRS *Z*-scores to be used directly in the conventional fine-mapping framework. We use the approach in Yang et al.^35^ with these summary statistics at given quantile level *τ* to compute PIPs and construct credible sets.

We now show the detailed derivation of eq. (2). Define the projection matrix onto the column space of **C** as **P**_*C*_ = **C C**^⊤^**C** −1 **C**^⊤^, and let 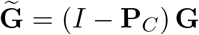 denote the matrix of covariate-residualized genotypes with *j*th column 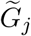. For SNP *j*, the QRS score statistic at quantile *τ* is

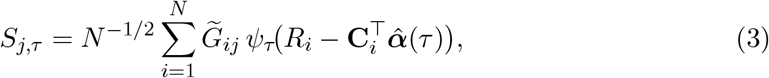

where *ψ*_*τ*_ (*u*) = *τ* − **1**(*u* < 0) and 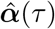 is the quantile regression estimate from the covariate-only model. Its variance is estimated by

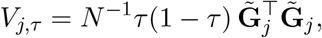

hence the standardized QRS *Z*-score is

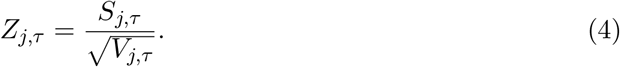

Let 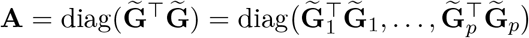. The diagonal variance matrix of the score statistics is then

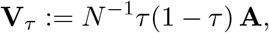

hence the standardized QRS *Z*-scores satisfy

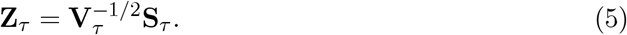

Standard Bahadur representation results for quantile regression^36;37^ imply that

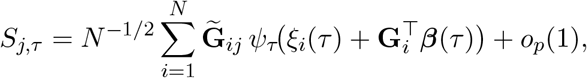

uniformly in *j*. In vector form,

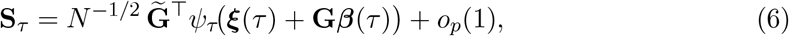

where *ψ*_*τ*_ is applied elementwise.

We now derive the expectation of **S**_*τ*_. Let *F*_*ξ*| **G**,**C**_ (· |**G**_*i*_, **C**_*i*_) be the conditional cdf of *ξ*_*i*_(*τ*) given (**G**_*i*_, **C**_*i*_), and define *f*_0_ := *f*_*ξ*|**G**,**C**_ (0 | **G**_*i*_, **C**_*i*_) to be the corresponding conditional density at 0. We assume *f*_0_ > 0 and that it is approximately constant near 0. Under the quantile model, *F*_*ξ*|**G**,**C**_ (0 | **G**_*i*_, **C**_*i*_) = *τ*. For a given individual *i*,

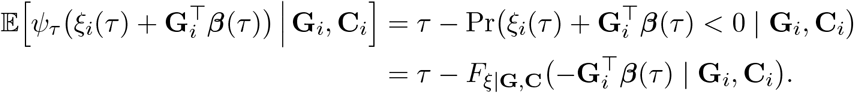

A first-order Taylor expansion of the cdf around 0 gives, for small local effects ***β***(*τ*),

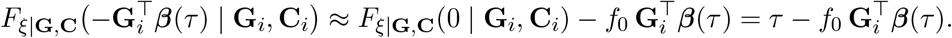

Hence

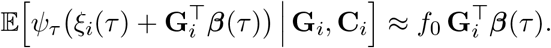

Plugging this into the expression for *S*_*j,τ*_ above, we get

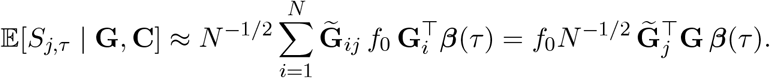

Because 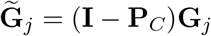 is orthogonal to the covariate space 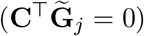, we have

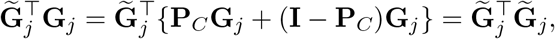

and in matrix form 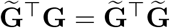. Hence

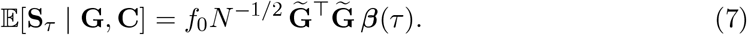

By the same arguments as in Wang et al.^9^, the covariance of the score vector satisfies

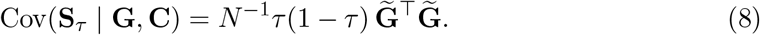

Using 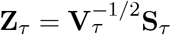 in eq. (5) with **V**_*τ*_ = *N* ^−1^*τ* (1 − *τ*)**A**, we obtain

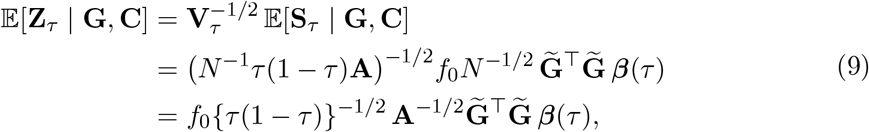

and

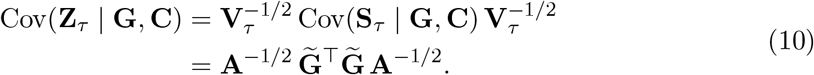

We define 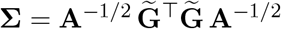, which is the *p* _×_ *p* correlation matrix of the residualized genotypes 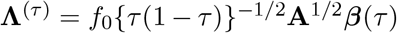. Let **Λ**^(*τ*)^ = *f*_0_ {*τ* (1 − *τ*)}^−1*/*2^**A**^1*/*2^***β***(*τ*) be the vector of scaled quantile effects. Then we can write

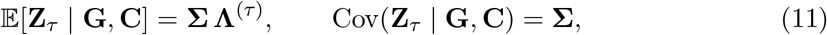

which is what we needed to show for eq. (2).

#### Credible sets

We define a credible set as the smallest subset of correlated variants (with correlation > *r*) that has probability *ρ* or greater of containing at least one causal variant.

## Data Availability

Processed data, including gene expression matrices and comprehensive molecular QTL summary statistics, can be accessed via the PigGTEx portal at http://piggtex.farmgtex.org.

## Code Availability

Regenie.QRS: https://github.com/FanWang0216/Regenie.QRS

Regenie: https://rgcgithub.github.io/Regenie/

CARMA: https://github.com/ZikunY/CARMA

g:Profiler: https://biit.cs.ut.ee/gprofiler/gost

AlphaGenome: https://deepmind.google.com/science/alphagenome/

MatchIt: https://github.com/kosukeimai/MatchIt

## Supplemental Figures and Tables

**Figure S1:**
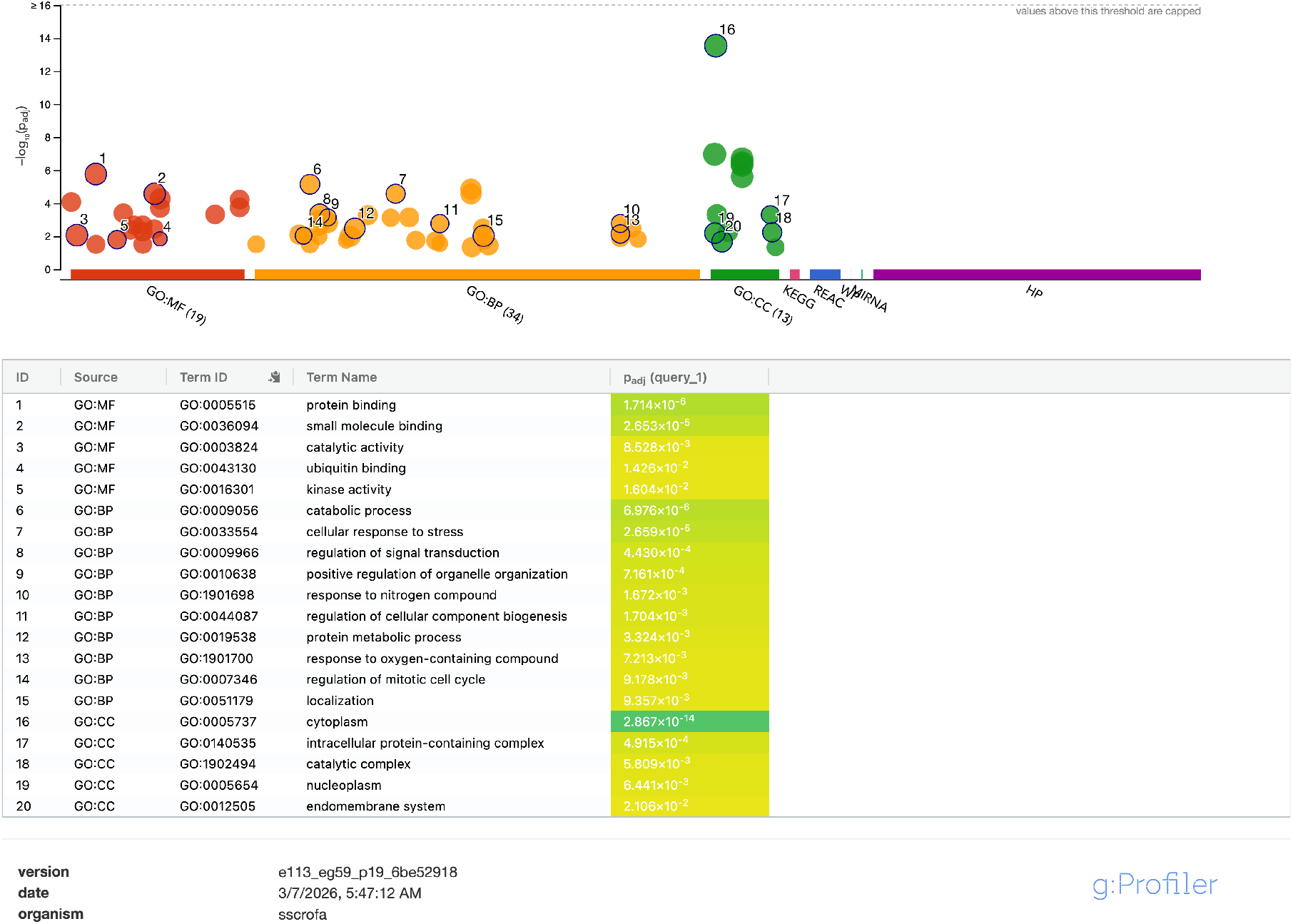
Functional enrichment analysis (g:Profiler) of Regenie.QRS only genes in blood. Manhattan plot of enriched pathways and terms across different molecular databases including Gene Ontology: Molecular Function (GO:MF), Biological Process (GO:BP), and Cellular Component (GO:CC), as well as KEGG, Reactome, and WikiPathways. Driver terms in each GO category are highlighted in the table below along with their adjusted p-values.

**Figure S2:**
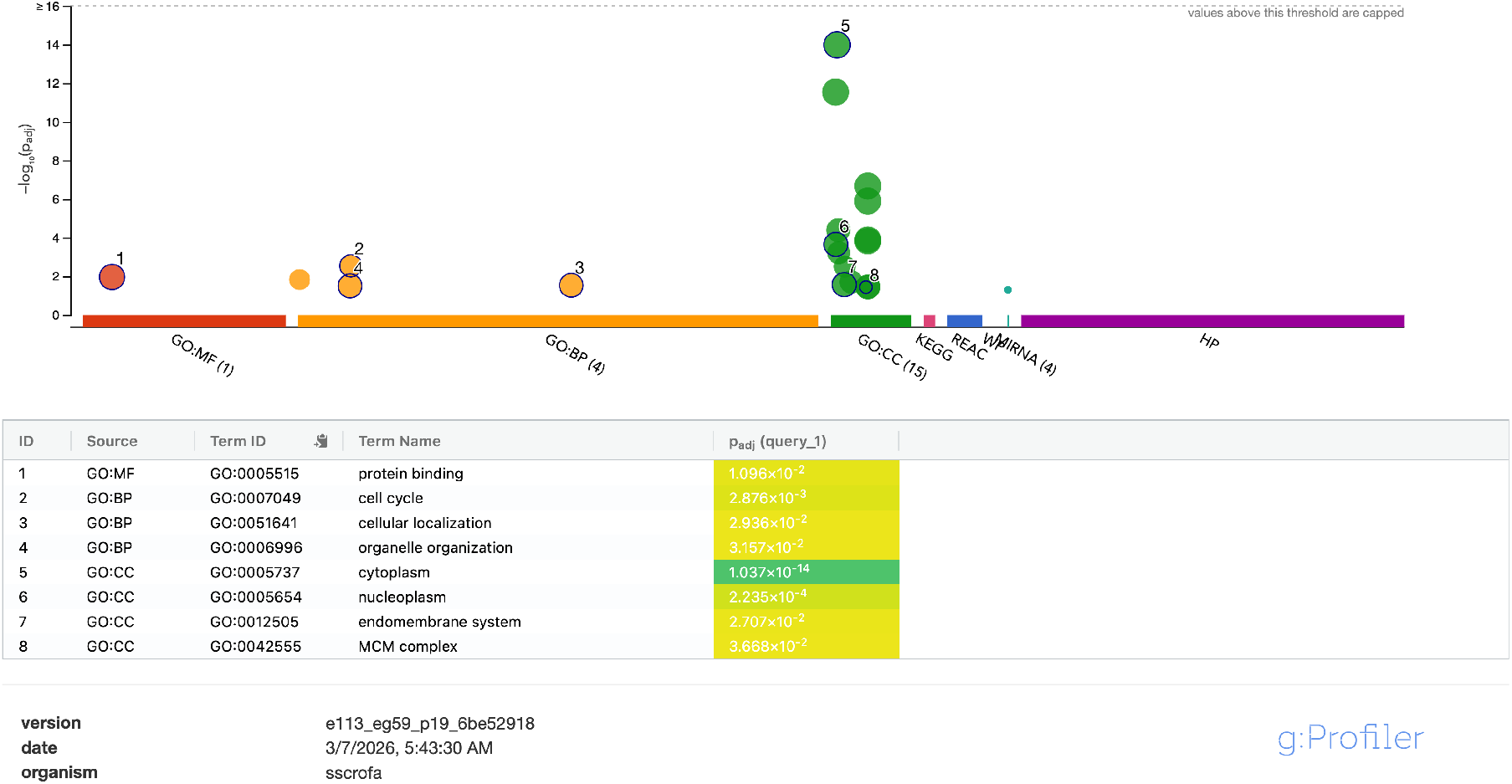
Functional enrichment analysis (g:Profiler) of Regenie only genes in blood. Manhattan plot of enriched pathways and terms across different molecular databases including Gene Ontology: Molecular Function (GO:MF), Biological Process (GO:BP), and Cellular Component (GO:CC), as well as KEGG, Reactome, and WikiPathways. Driver terms in each GO category are highlighted in the table below along with their adjusted p-values.

**Figure S3:**
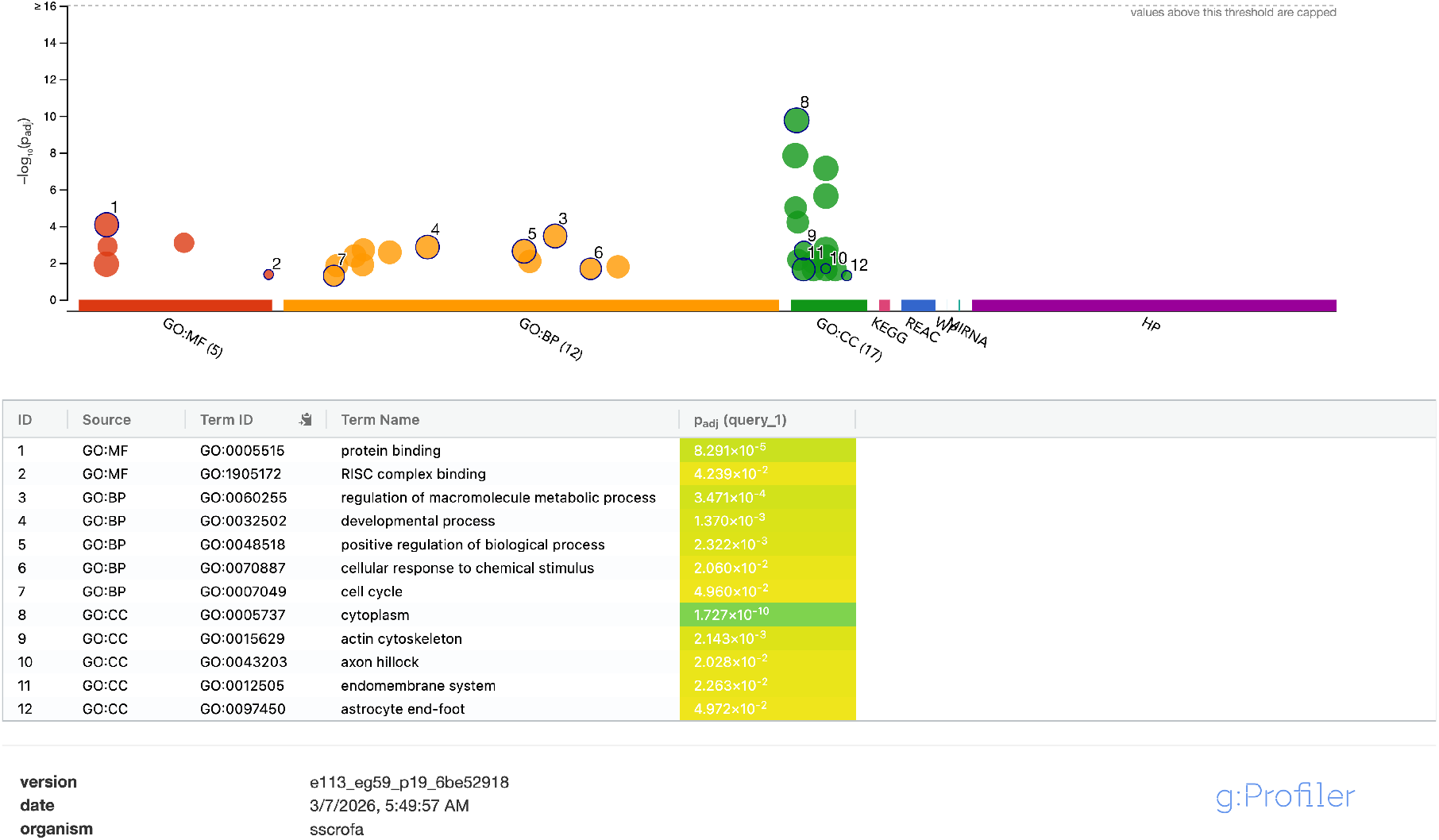
Functional enrichment analysis (g:Profiler) of Regenie.QRS only genes in brain. Manhattan plot of enriched pathways and terms across different molecular databases including Gene Ontology: Molecular Function (GO:MF), Biological Process (GO:BP), and Cellular Component (GO:CC), as well as KEGG, Reactome, and WikiPathways. Driver terms in each GO category are highlighted in the table below along with their adjusted p-values.

**Figure S4:**
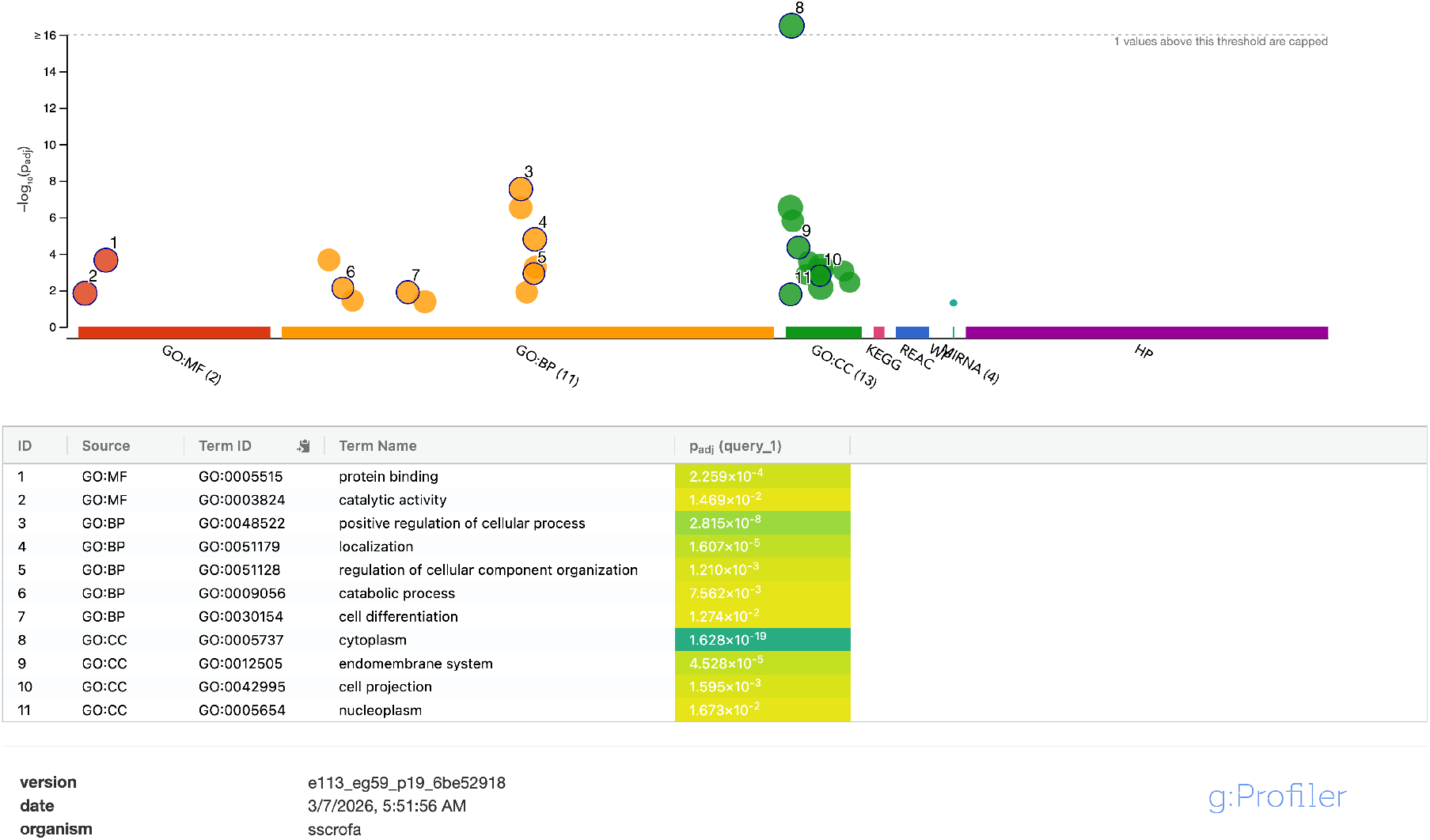
Functional enrichment analysis (g:Profiler) of Regenie only genes in brain. Manhattan plot of enriched pathways and terms across different molecular databases including Gene Ontology: Molecular Function (GO:MF), Biological Process (GO:BP), and Cellular Component (GO:CC), as well as KEGG, Reactome, and WikiPathways. Driver terms in each GO category are highlighted in the table below along with their adjusted p-values.

**Figure S5:**
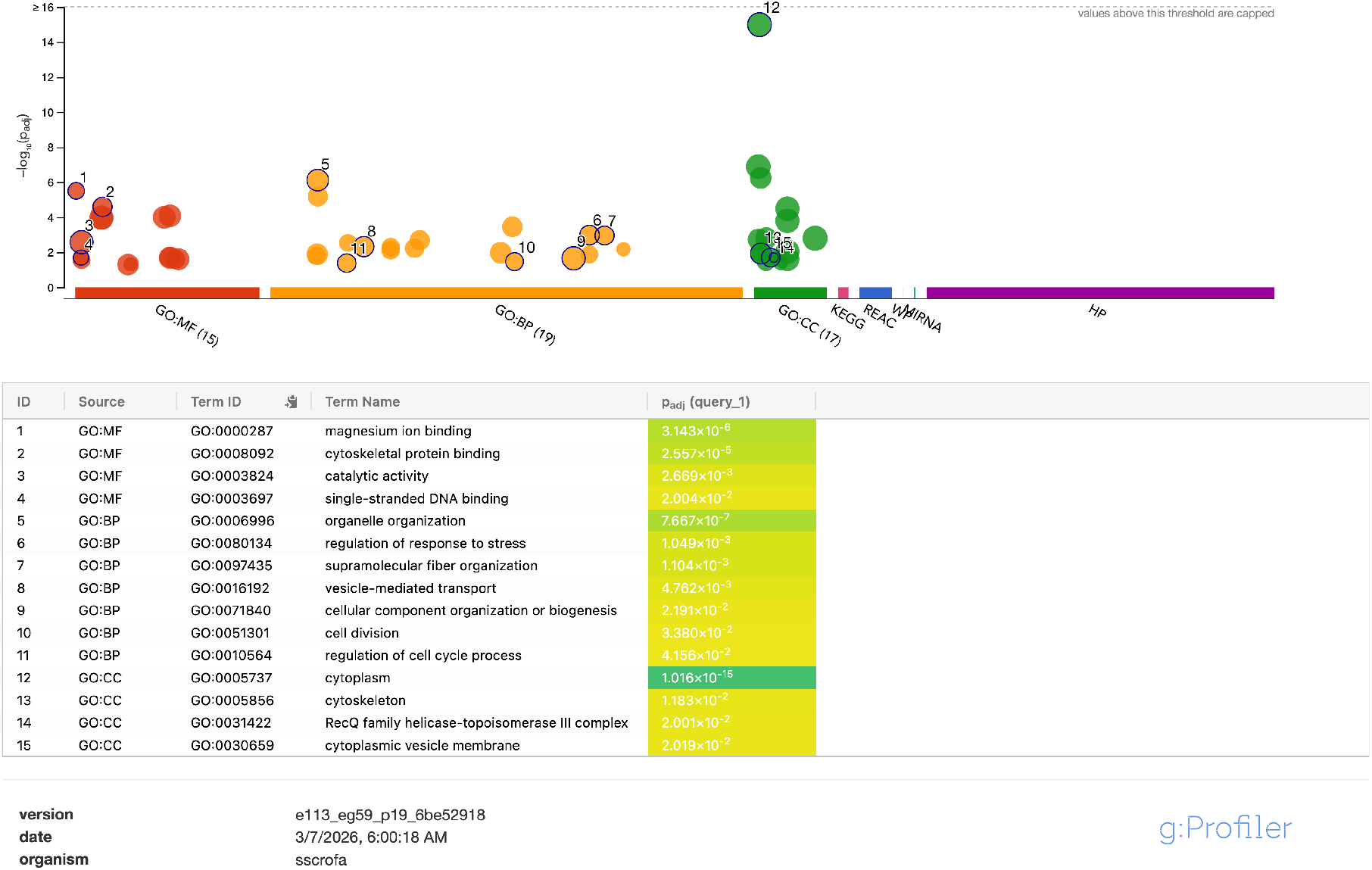
Functional enrichment analysis (g:Profiler) of Regenie.QRS only genes in adipose. Manhattan plot of enriched pathways and terms across different molecular databases including Gene Ontology: Molecular Function (GO:MF), Biological Process (GO:BP), and Cellular Component (GO:CC), as well as KEGG, Reactome, and WikiPathways. Driver terms in each GO category are highlighted in the table below along with their adjusted p-values.

**Figure S6:**
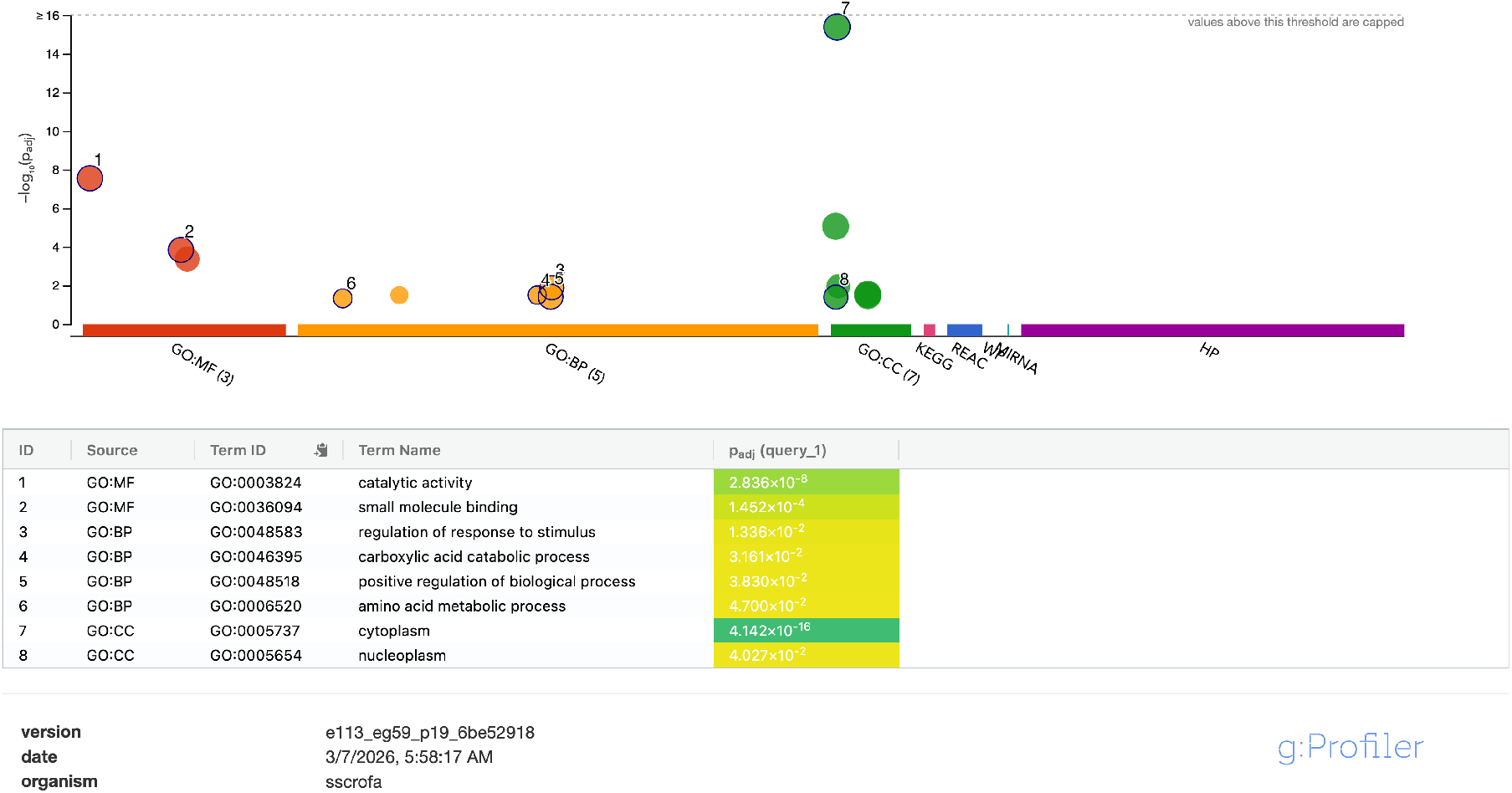
Functional enrichment analysis (g:Profiler) of Regenie only genes in adipose. Manhattan plot of enriched pathways and terms across different molecular databases including Gene Ontology: Molecular Function (GO:MF), Biological Process (GO:BP), and Cellular Component (GO:CC), as well as KEGG, Reactome, and WikiPathways. Driver terms in each GO category are highlighted in the table below along with their adjusted p-values.

**Figure S7:**
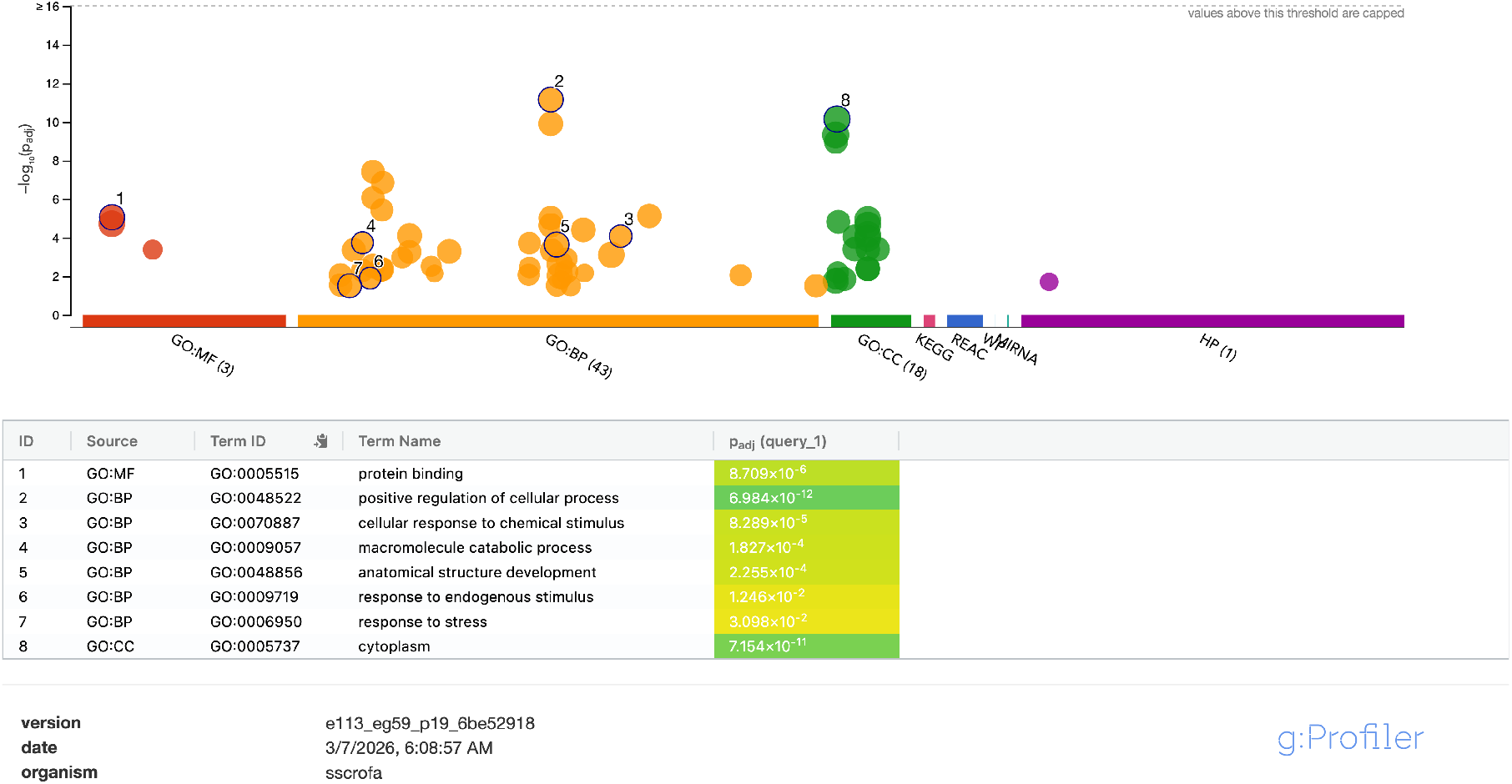
Functional enrichment analysis (g:Profiler) of Regenie.QRS only genes in muscle. Manhattan plot of enriched pathways and terms across different molecular databases including Gene Ontology: Molecular Function (GO:MF), Biological Process (GO:BP), and Cellular Component (GO:CC), as well as KEGG, Reactome, and WikiPathways. Driver terms in each GO category are highlighted in the table below along with their adjusted p-values.

**Figure S8:**
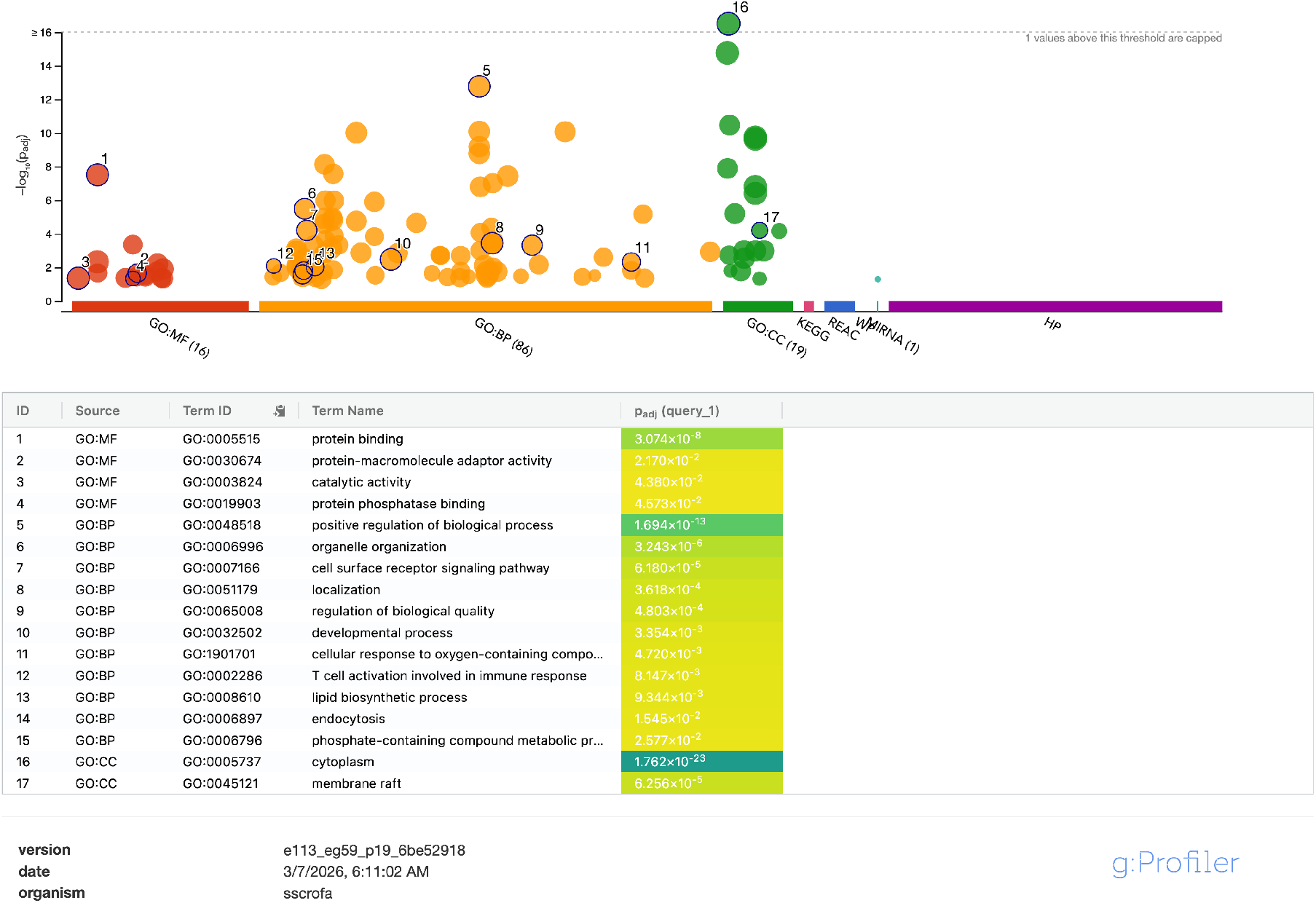
Functional enrichment analysis (g:Profiler) of Regenie only genes in muscle. Manhattan plot of enriched pathways and terms across different molecular databases including Gene Ontology: Molecular Function (GO:MF), Biological Process (GO:BP), and Cellular Component (GO:CC), as well as KEGG, Reactome, and WikiPathways. Driver terms in each GO category are highlighted in the table below along with their adjusted p-values.

**Figure S9:**
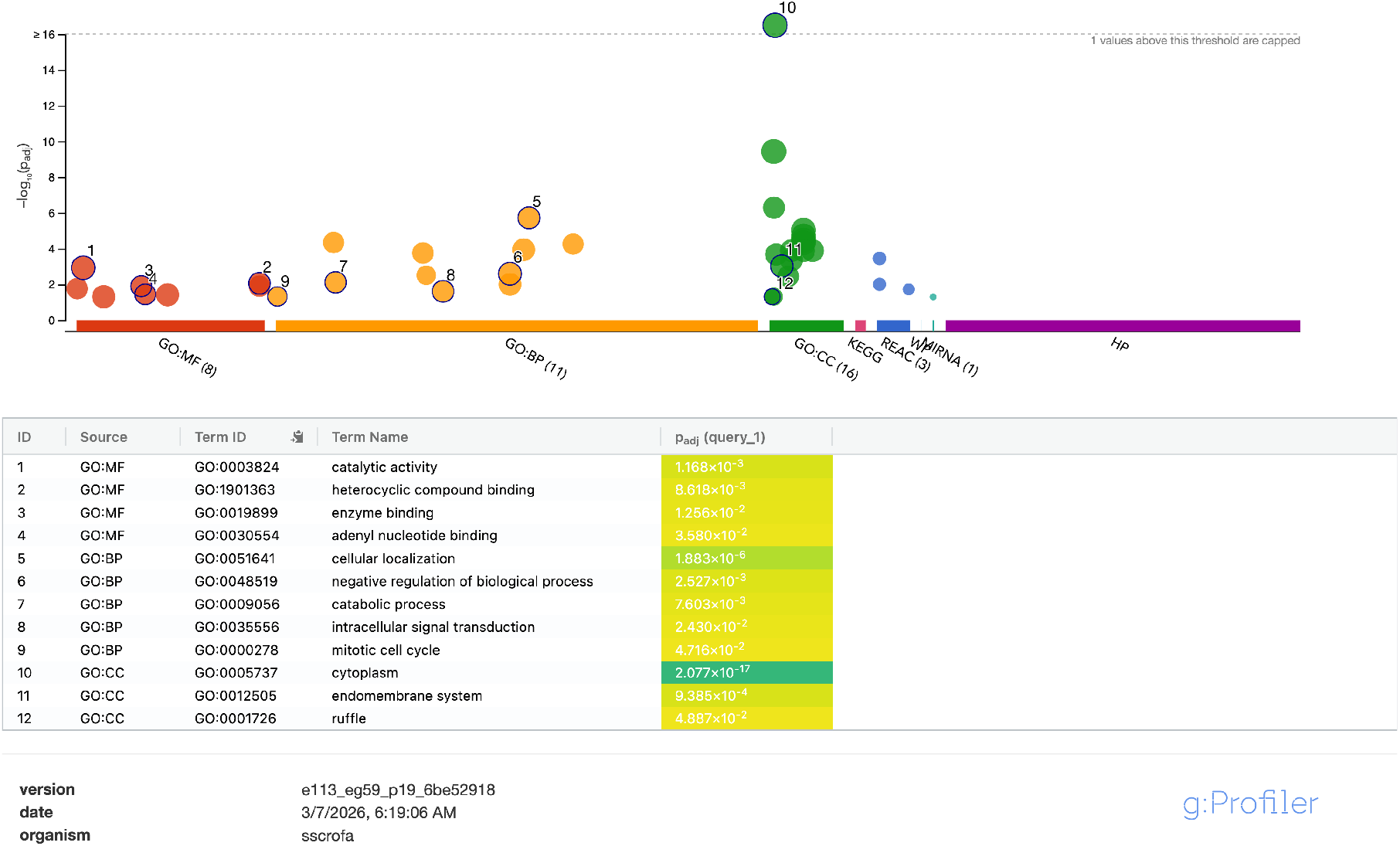
Functional enrichment analysis (g:Profiler) of Regenie.QRS only genes in liver. Manhattan plot of enriched pathways and terms across different molecular databases including Gene Ontology: Molecular Function (GO:MF), Biological Process (GO:BP), and Cellular Component (GO:CC), as well as KEGG, Reactome, and WikiPathways. Driver terms in each GO category are highlighted in the table below along with their adjusted p-values.

**Figure S10:**
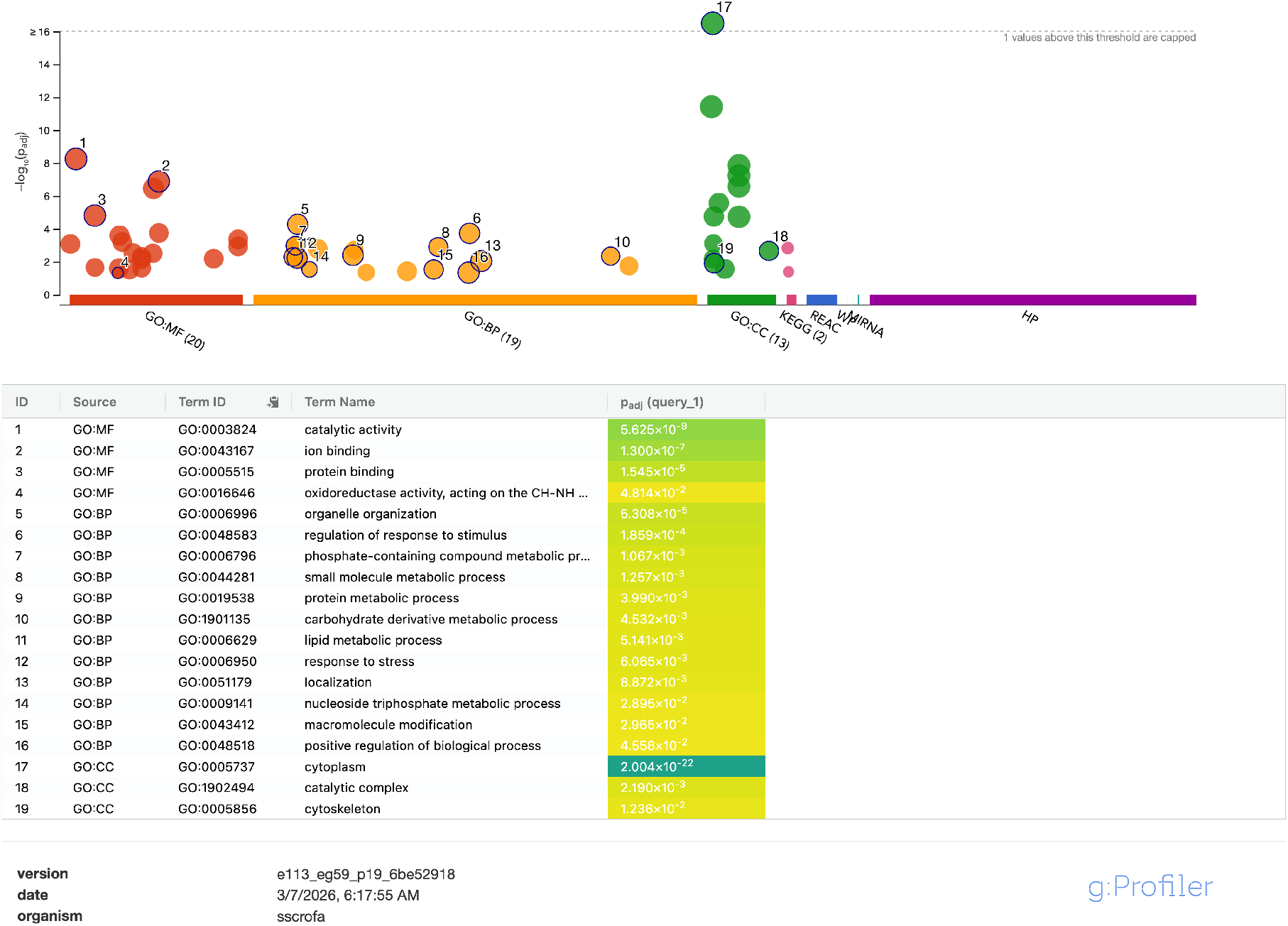
Functional enrichment analysis (g:Profiler) of Regenie only genes in liver. Manhattan plot of enriched pathways and terms across different molecular databases including Gene Ontology: Molecular Function (GO:MF), Biological Process (GO:BP), and Cellular Component (GO:CC), as well as KEGG, Reactome, and WikiPathways. Driver terms in each GO category are highlighted in the table below along with their adjusted p-values.

**Figure S11:**
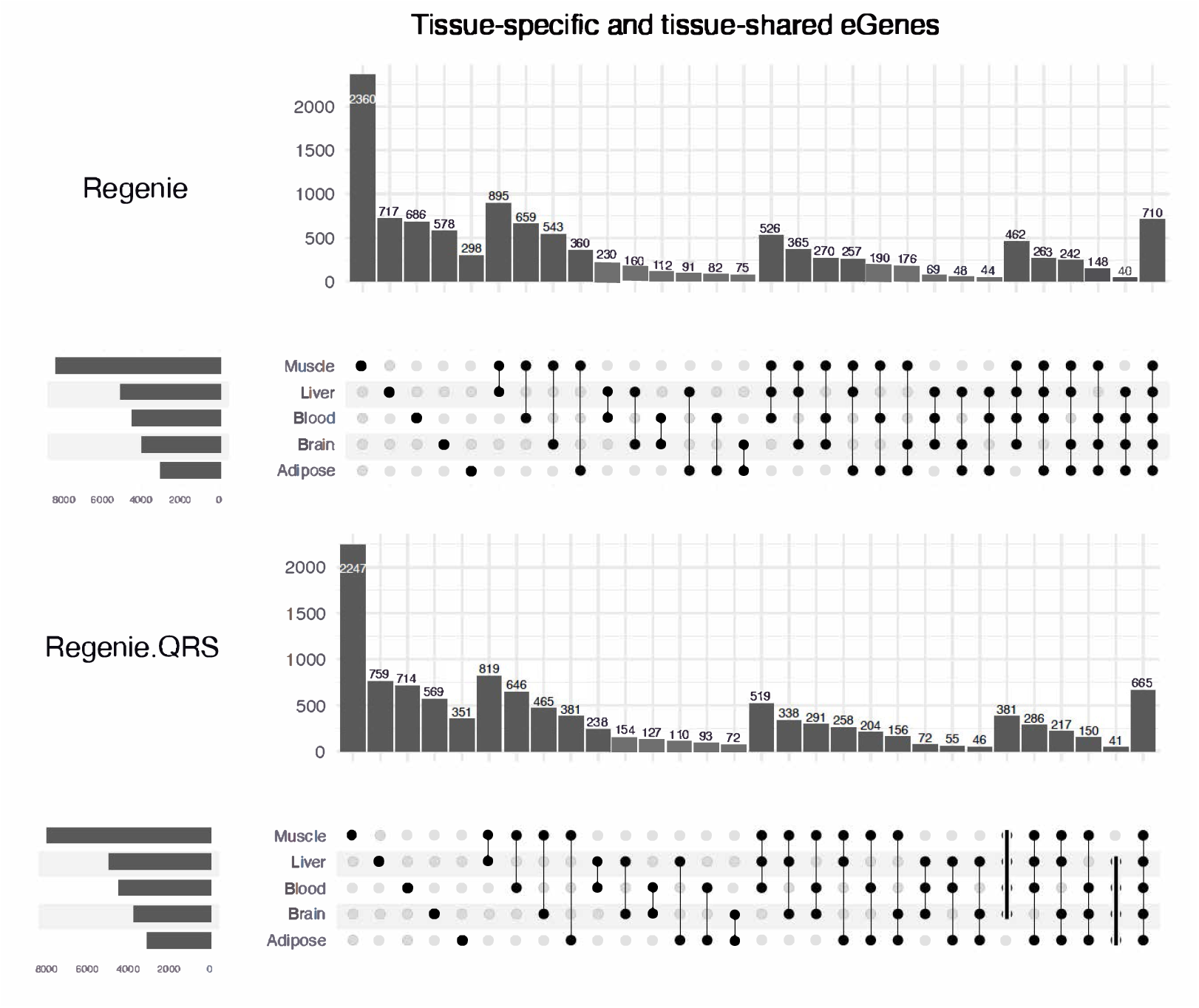
Tissue-specific and shared eGenes identified by Regenie and Regenie.QRS across five tissues. Left bars indicate the number of eGenes per tissue; top bars show intersection sizes for the 30 most shared tissue combinations. Black dots denote tissues included in each intersection.

**Figure S12:**
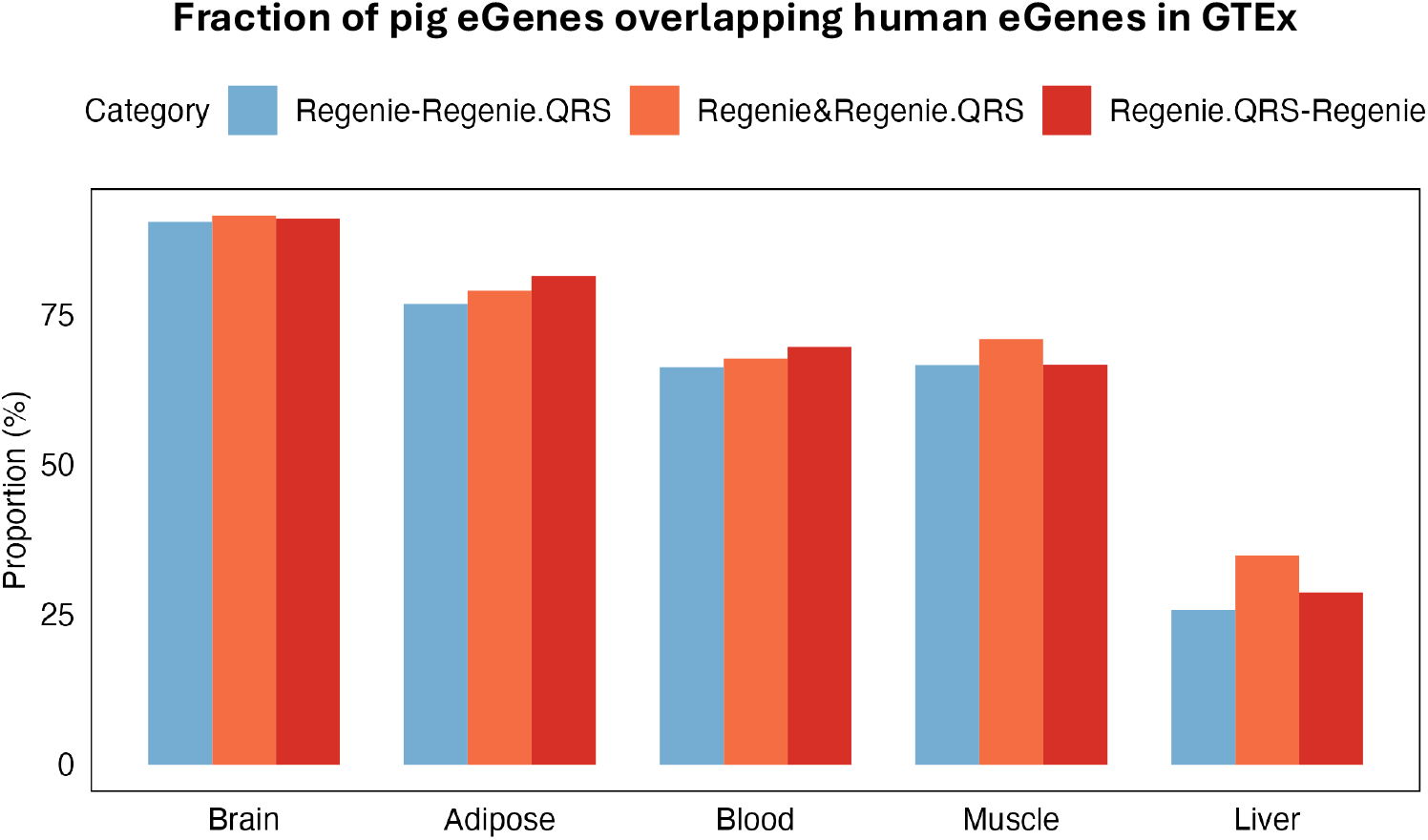
eGenes sharing between pigs and humans in matching tissues. Shown are the proportions of pig eGenes that are also reported as eGenes in human GTEx.

**Figure S13:**
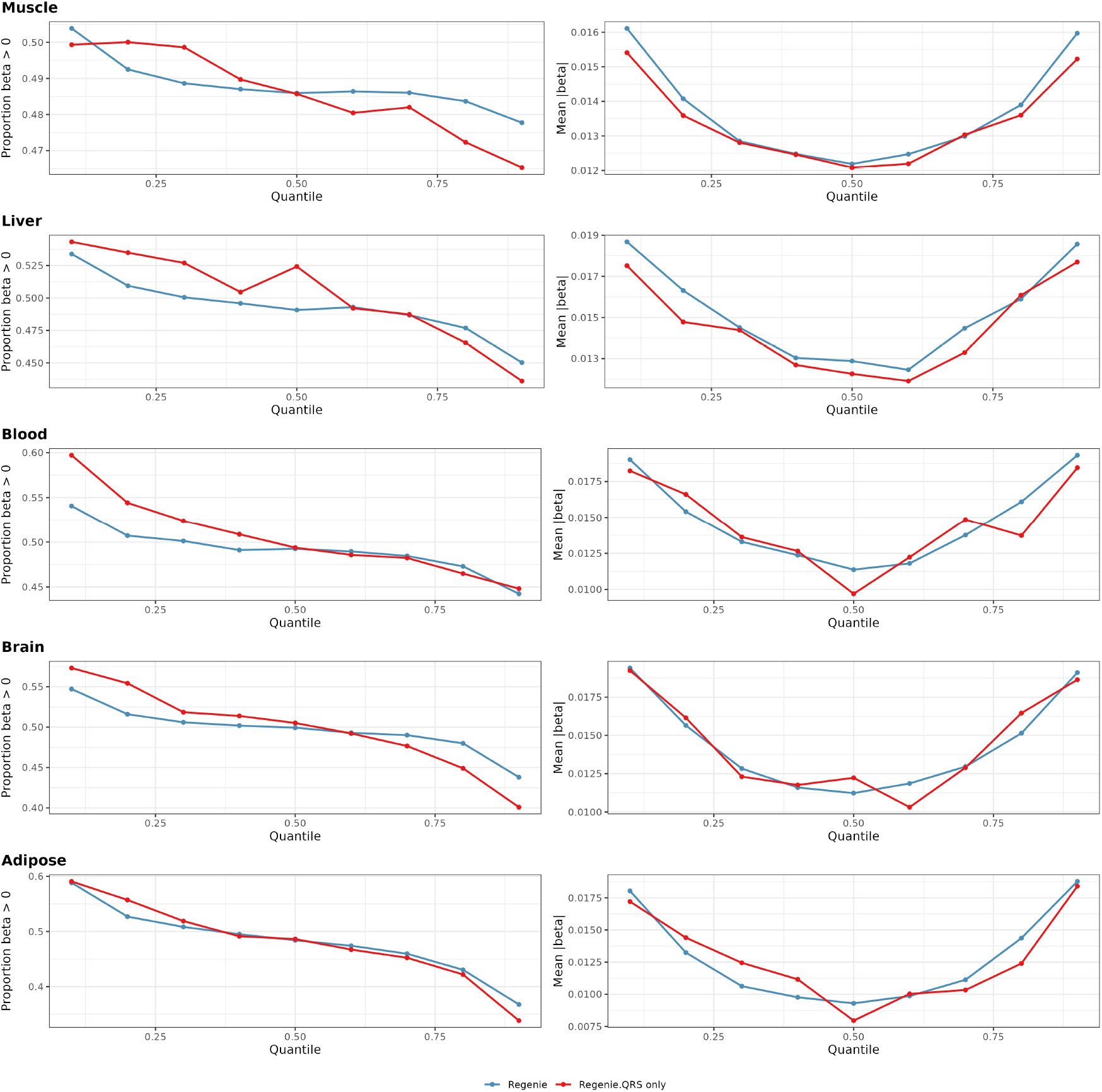
Quantile-dependent genetic effects and directional bias in PigG-TEx data. Results are shown for eQTLs identified by different methods: Regenie and Regenie.QRS only. **Left**. Proportion of positive beta coefficients (*β* > 0) across quantiles. **Right**. Mean absolute beta values across quantiles (*τ*) exhibiting a U-shaped pattern, indicating larger effect sizes at the distributional tails.

**Figure S14:**
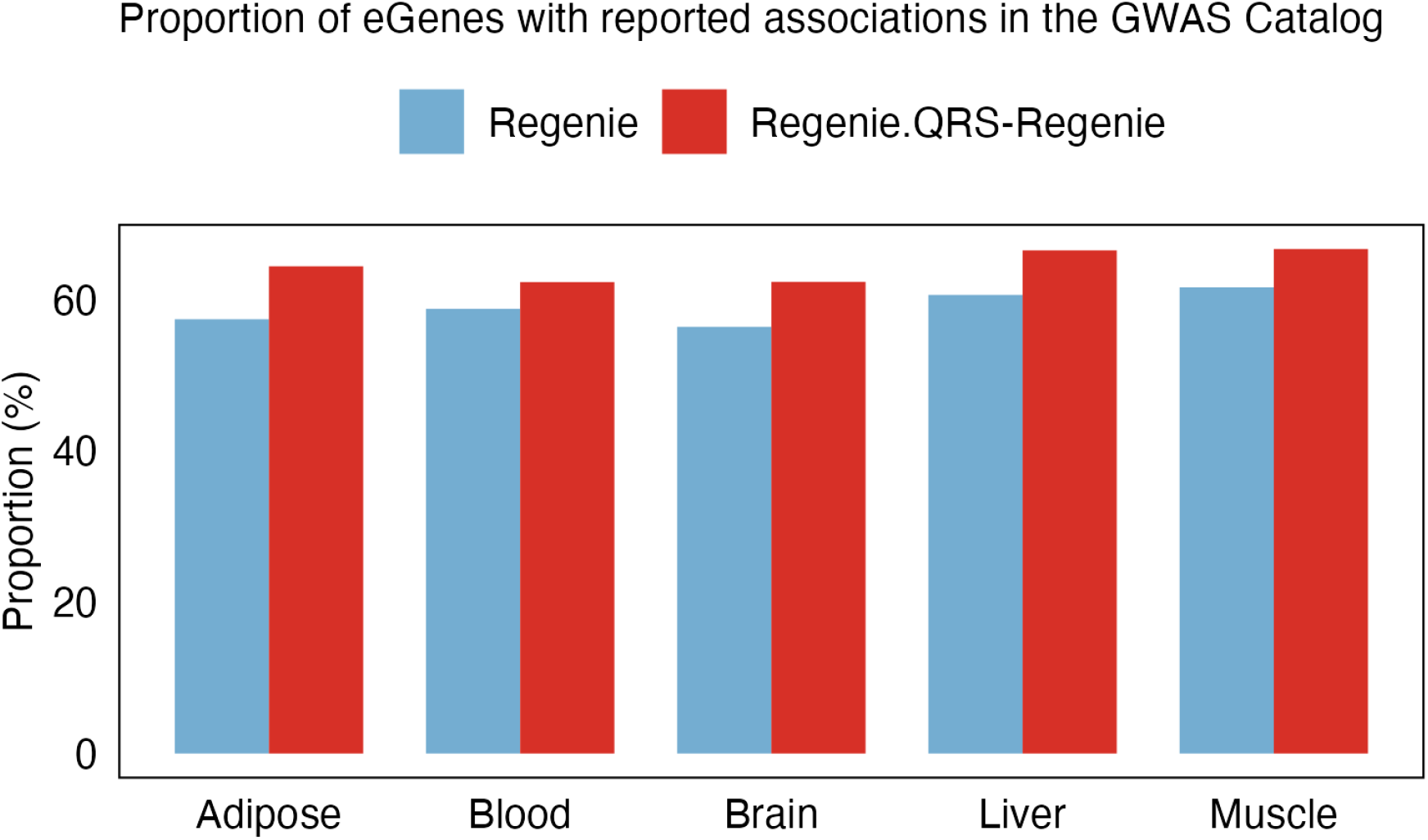
Proportion of eGenes with reported associations in the GWAS Catalog across tissues. For each tissue, the percentage of pig eGenes identified using Regenie and Regenie.QRS only that have known associations in the GWAS Catalog are reported.

**Figure S15:**
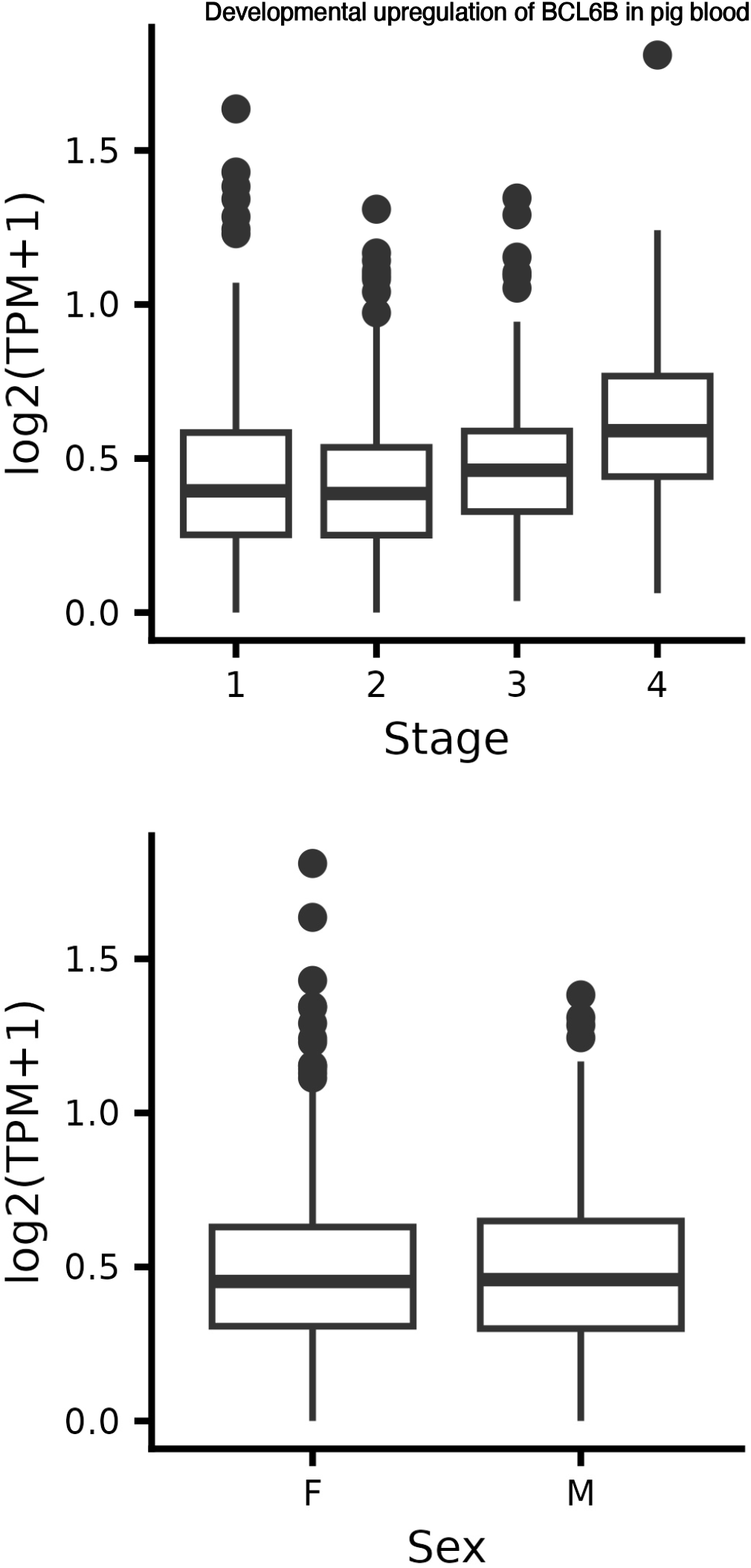
BCL6B expression levels by developmental stage and sex in the Pig developmental GTEx project. Each stage includes ~ 150 pigs, with samples being independent of the PigGTEx data analyzed elsewhere in the paper.

**Figure S16:**
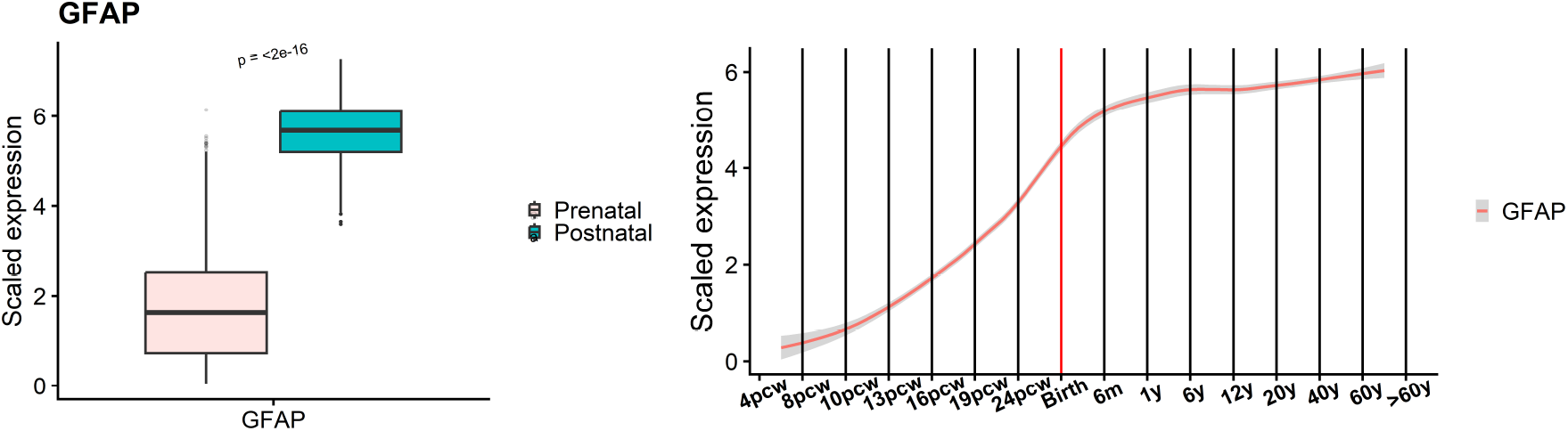
Temporal expression dynamics of *GFAP* across developmental stages^30^. **Left**. Boxplots comparing scaled *GFAP* expression between prenatal and postnatal samples. **Right**. Longitudinal trajectory of scaled expression across the lifespan (years).

**Table S1:** AlphaGenome molecular modalities used to predict variant effects.

| Modality | Description |
| --- | --- |
| RNA expression (RNA-seq) | Steady-state transcript levels across tissues |
| Transcription initiation (CAGE) | Activity at transcription start sites |
| Transcription initiation (PRO-cap) | Nascent capped RNA initiation signal |
| Splice sites | Locations where splicing begins or ends |
| Splice site usage | Quantitative usage of splice sites |
| Splice junctions | Counts and strength of exon–exon junctions |
| Chromatin accessibility (DNase-seq) | Open chromatin regions |
| Chromatin accessibility (ATAC-seq) | Alternative measure of open chromatin |
| Histone modifications (ChIP-seq) | Chromatin state marks |
| Transcription factor binding (ChIP-TF) | Binding of regulatory proteins |
| Chromatin contact maps (Hi-C/Micro-C) | 3D genome architecture |

**Table S2:**
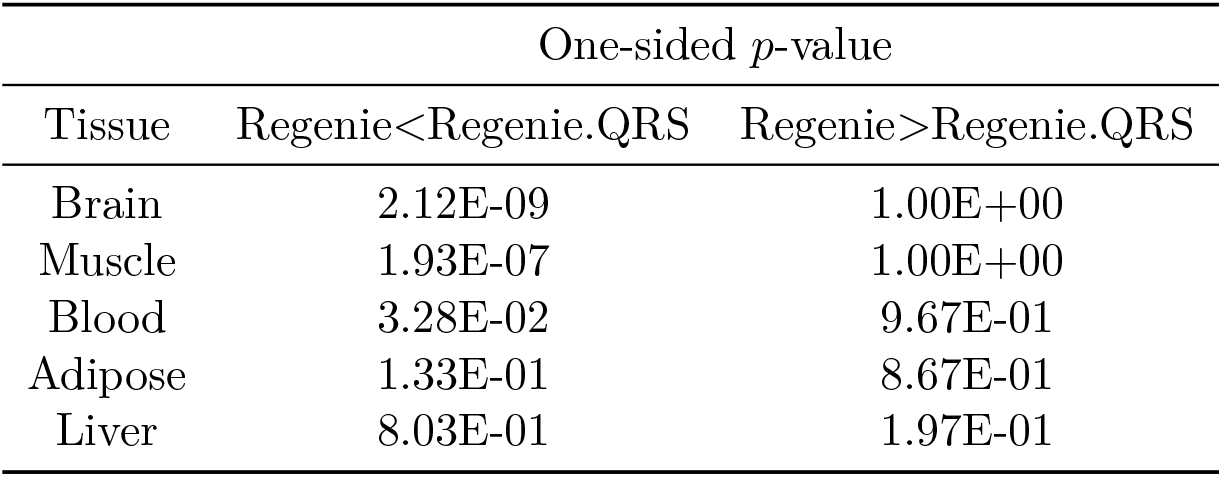
Tissue-matched regulatory effects independent of target gene. Comparison of AlphaGenome scores (RNA-seq) for eQTLs identified by Regenie and Regenie.QRS. *p*-values are from one-sided Wilcoxon test.

**Table S3:** Tissue- and gene-matched regulatory effects. Comparison of AlphaGenome scores (RNA-seq) for eQTLs identified by Regenie and Regenie.QRS. *p*-values are from one-sided Wilcoxon test.

| Tissue | One-sided $p$-value | |
| --- | --- | --- |
|  | Regenie<Regenie.QRS | Regenie>Regenie.QRS |
| Brain | 2.64E-09 | 1.00E+00 |
| Blood | 1.00E+00 | 2.74E-09 |
| Adipose | 1.00E+00 | 3.08E-04 |
| Muscle | 7.65E-02 | 9.23E-01 |
| Liver | 6.51E-01 | 3.49E-01 |

## Notes

### Competing Interest Statement

The authors have declared no competing interest.

